# Anticipatory coadaptation of ankle stiffness and sensorimotor gain for standing balance

**DOI:** 10.1101/506493

**Authors:** Charlotte Le Mouel, Romain Brette

**Affiliations:** Max Planck Institute for Intelligent Systems, Stuttgart, Germany; Sorbonne Université, INSERM, CNRS, Institut de la Vision, 17 rue Moreau, F-75012 Paris, France

## Abstract

External perturbation forces may compromise standing balance. The nervous system can intervene only after a delay greater than 100 ms, during which the body falls freely. With ageing, sensorimotor delays are prolonged, posing a critical threat to balance. We study a generic model of stabilisation with neural delays to understand how the organism should adapt to challenging balance conditions. The model suggests that ankle stiffness should be increased in anticipation of perturbations, for example by muscle co-contraction, so as to slow down body fall during the neural response delay. Increased ankle muscle co-contraction is indeed observed in young adults when standing in challenging balance conditions, and in older relative to young adults during normal stance. In parallel, the analysis of the model shows that increases in either stiffness or neural delay must be coordinated with decreases in spinal sensorimotor gains, otherwise the feedback itself becomes destabilizing. Accordingly, a decrease in spinal feedback is observed in challenging conditions, and with age-related increases in neural delay. These observations have been previously interpreted as indicating an increased reliance on cortical rather than spinal control of balance, despite the fact that cortical responses have a longer latency. Our analysis challenges this interpretation by showing that these observations are consistent with a functional coadaptation of spinal feedback gains to functional changes in stiffness and neural delay.

**Author summary:** Being able to stand still can be difficult when faced with an unexpected push. It takes the nervous system more than a tenth of a second to respond to such a perturbation, and during this delay the body falls under the influence of its own weight. By co-contracting their ankle muscles in anticipation of a perturbation, subjects can increase their ankle stiffness, which slows down their fall during the neural delay. Young subjects indeed adopt this strategy when they need to remain particularly still (for example when they stand in front of a cliff). Older subjects adopt this strategy even during normal standing. We present a model of standing balance that shows that this postural strategy provides partial compensation for the increase in neural delays with ageing. According to our model, increasing ankle stiffness only improves balance if it is accompanied by a decrease in sensorimotor gain. This provides a novel and functional interpretation for the decrease in spinal feedback observed during ageing, and observed in young subjects when they stand in challenging balance conditions.

## I. Introduction

External perturbation forces may compromise standing balance (1). Current theories emphasise the role of the motor cortex in the feedback correction of unexpected perturbations (2,3). Indeed, the spinal feedback correction of perturbations is reduced when standing in challenging balance conditions (Fig 1.A, B), such as when standing facing a cliff (4), when standing on a narrow support (5) or simply when closing the eyes (6). The classical interpretation for this reduction in the spinal contribution to balance is that, in challenging conditions, the control of balance is delegated to supra-spinal structures, such as the cortex, which may allow for a more refined control than the spinal cord (3). Even in normal standing conditions, the spinal feedback correction of perturbations is reduced in older adults relative to young adults (Fig 1.C) (7–9), as well as delayed (10,11). The interpretation for this is likewise that older subjects rely more on cortical rather than spinal control of balance (2).

**Fig 1.**
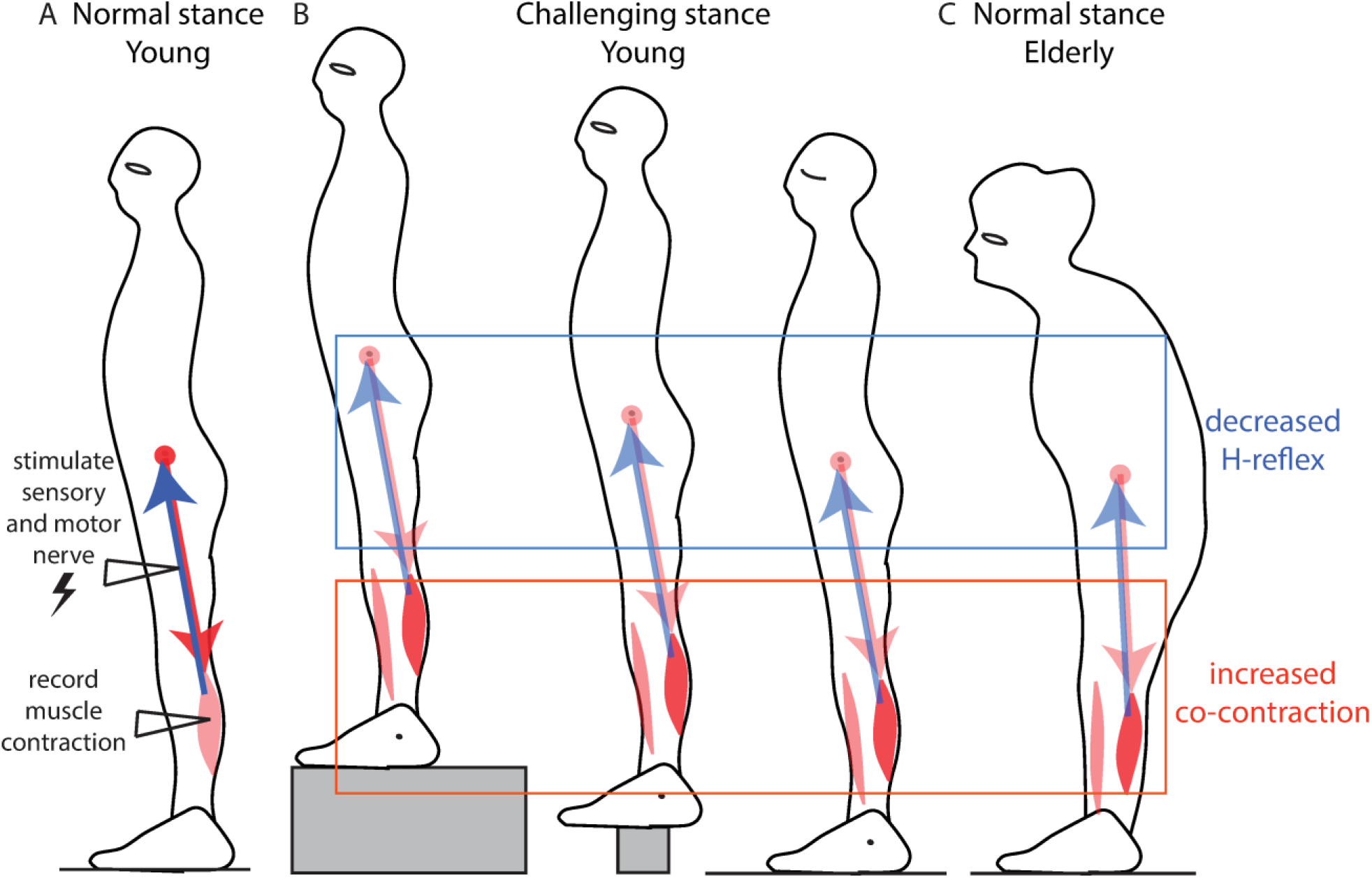
Ankle stiffness and spinal feedback. A. Spinal feedback is probed by electrically stimulating the sensory fibres of a muscle and measuring the resulting change in muscle contraction, called the H-reflex. B. Decrease in soleus H-reflex in challenging balance conditions: when standing facing a cliff (left) (4), when standing on a narrow support (middle) (5) and standing with the eyes closed (right) (6). Co-contraction of antagonist ankle muscles is observed in each of these three cases. C. Decrease in soleus H-reflex in older adults in normal stance (7–9). Co-contraction of antagonist ankle muscles during quiet standing is observed in older but not in young subjects (7,12).

The difficulty with this approach is that the neural feedback control of movement introduces delays. The shortest spinal delay after which a change in contraction can be observed following a direct electrical stimulation of the sensory nerve (as in Fig 1) is around 30 ms in young subjects (10,11). However, after a perturbation of stance, the earliest change in muscular contraction occurs with a longer delay of around 100 ms in young subjects (13,14). In older subjects, this is further delayed by 10 to 30 ms (13,14). The change in force due to this muscle contraction is only observed after an additional 40 ms (15). Responses involving the cortex have even longer delays than spinal feedback (16). It is well known from control theory that delays are critical when using sensory feedback to counteract external perturbations (17): thus, a system that is stabilized by feedback control may become unstable simply if the control delay increases. Faster contributions to balance may therefore be more effective than cortical control, particularly when standing in challenging balance conditions, or with age-related increases in neural delays.

During this neural response delay after a perturbation, the movement of the body is entirely determined by the body and environmental mechanical properties, such as stiffness, inertia and weight. Ankle stiffness can be actively increased through co-contraction of the ankle muscles (18). Young subjects co-contract the ankle muscles when standing in challenging balance conditions (5,6,19), and older subjects co-contract the ankle muscles even in normal standing conditions (Fig 1.C)(7,12).

To understand the relative contributions of ankle stiffness, neural delay and sensorimotor gain to balance performance, we analyze a widely used model of standing balance. We propose a novel method to determine how sensorimotor gains should adjust to changes in ankle stiffness and neural delay. We show that the most effective strategy to improve balance relies on the co-adaptation of ankle stiffness and sensorimotor gains. Increasing ankle stiffness in advance of a perturbation can improve robustness to perturbations, by reducing the amplification of perturbations during the neural feedback delay. However, this improves balance only if the increase in ankle stiffness is coordinated with a decrease in sensorimotor gain, to prevent over-compensation. This strategy is also the most effective for maintaining balance performance despite increasing neural delay.

The decreased spinal feedback observed during normal standing in older subjects, and during standing in challenging balance conditions in young subjects, should therefore not be interpreted as the mark of a more cortical control of balance, but on the contrary as the sign that motor stabilisation is achieved through an increased reliance on ankle stiffness.

## II. Modelling results

After a perturbation of stance, the earliest change in muscular contraction occurs with a delay of around 100 ms in young subjects (13,14). The change in force due to this muscle contraction is only observed after an additional 40 ms (15). During this neural response delay *τ_delay_* = 0.14 *s*, the body centre of mass is accelerated by gravity. We model the body mechanics using the single inverted pendulum model, which is widely used to model human stance in the sagittal plane (20,21). The ankle angle *θ* follows:

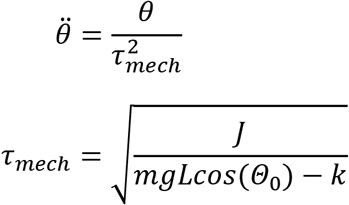

The dynamics are thus governed by the mechanical time constant *τ_mech_*, which captures the effects of the rotational inertia around the ankles *J*, ankle stiffness *k* and weight *mg*. Any increase in ankle stiffness *k* up to the critical ankle stiffness *K_crit_* = *mgLcos*(*Θ*_0_) increases the mechanical time constant of the body. Ankle stiffness thus slows down falling during the response delay (22). During this delay, perturbations are amplified by a factor 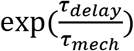, as shown in the Methods. This suggests that one strategy to improve immobility in challenging balance conditions may be to increase ankle stiffness so as to decrease the relative speed 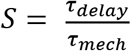. Increasing ankle stiffness may likewise help to compensate for an increase in feedback delay, by mitigating the increase in *S*.

We wish to determine how the neural feedback control of balance is affected by an increase in ankle stiffness, with and without an increase in neural delay. For this, we first assume a simple model for the control of standing balance, and show that both strategies require a decrease in sensorimotor gain. We then show that these results generalise to more complex neuro-mechanical models.

We first assume that balance is achieved through delayed feedback control of position and speed. Such a control strategy has previously been suggested to model human balance in both the sagittal (21) and lateral (23) plane. This delayed proportional-derivative control has the form:

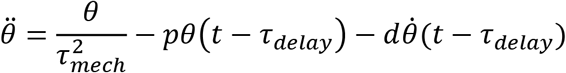

### 1. Stability limits

For any combination of mechanical parameters (determining *τ_mech_*) and neural response delay, there is a limited range of proportional gains *p* and derivative gains *d* that can stabilize the system (21). The equations of the stability boundaries are derived in the Supplementary (I) and provided in the Methods. They are plotted for increasing stiffness and neural delay in Fig 2.

**Fig 2.**
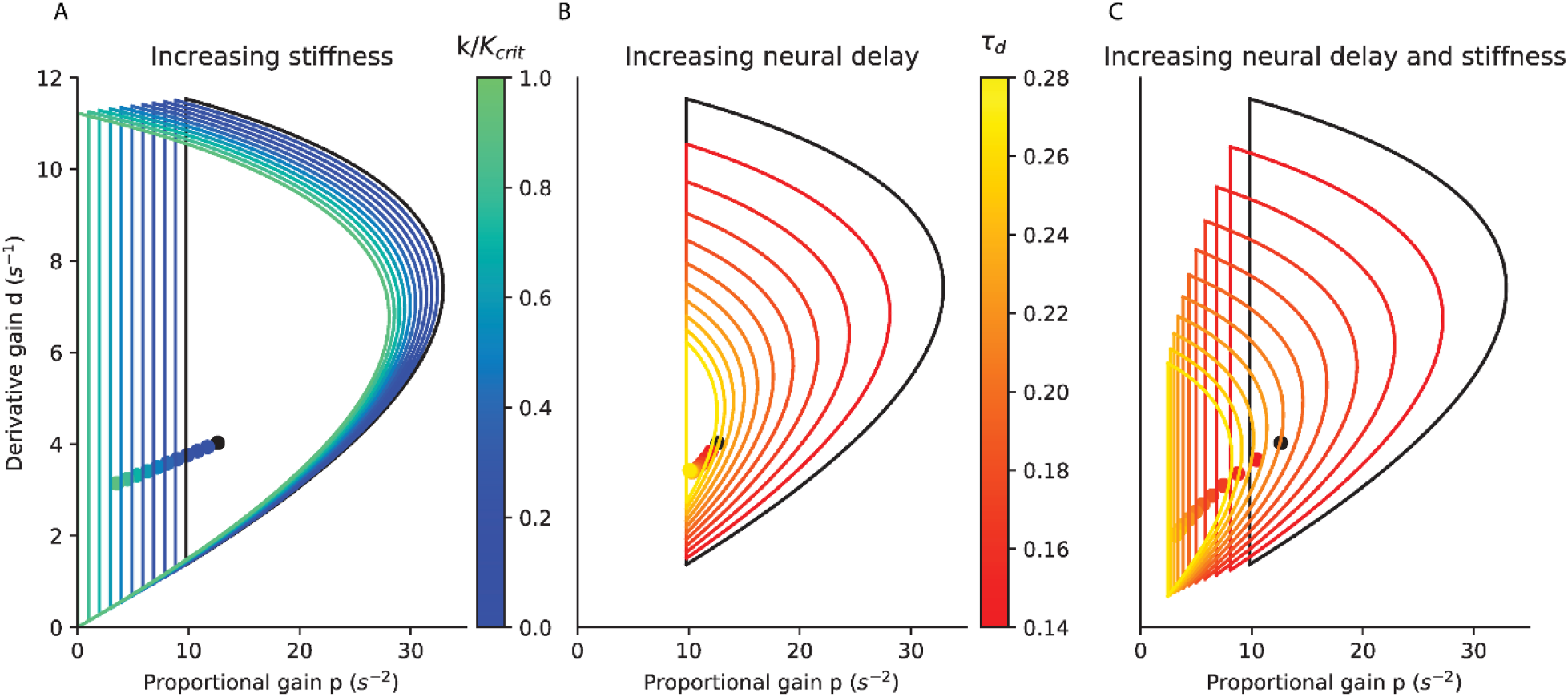
Stability limits of the proportional and derivative gains. A. for stiffness increasing from 0 (black) to critical stiffness (turquoise) with a delay of 0.14 s; B. for a delay increasing from 0.14 s (black) to 0.28 s (yellow) without stiffness; C. for combined increases in delay and stiffness (maintaining a constant S), from a delay of 0.14 s with a stiffness of 0 (black) to a delay of 0.28 s with a stiffness of 0.75 K_crit_ (yellow). The critical feedback gains for each condition are indicated as dots. In the three panels, the black curves are identical and correspond to zero stiffness and a delay of 0.14 s.

With increasing ankle stiffness (i.e. increasing *τ_mech_*, Fig 2.A), the minimal and maximal gains shift towards smaller values. Large values of feedback gains which are stable without ankle stiffness (Fig 2.A, black) become unstable with critical ankle stiffness (Fig 2.A, turquoise). Small values of feedback gains which are unstable without ankle stiffness become stable with critical ankle stiffness. This suggests that, if subjects increase ankle stiffness they may need to decrease their sensorimotor gains.

With aging, there is in increase in feedback delay (13,14). According to our model, with increasing feedback delay (Fig 2.B), the range of stable feedback gain decreases. The maximal gains decrease sharply. If the subject increases ankle stiffness to compensate for the increasing delay (for example by keeping *S* constant, Fig 2.C), then the effects of increasing stiffness (Fig 2. A) and delay (Fig 2.B) combine. The proportional gains which can stabilise the system for *τ_deiay_* = 0.14 *s* without ankle stiffness (Fig 2.C, black) are thus too large to stabilise the system after a doubling of the delay (Fig 2.C, yellow, *τ_delay_* = 0.28 *s, k* = 0.75 *K_crit_*). This suggests that when subjects increase ankle stiffness to compensate for an increase in neural delay, they must decrease their sensorimotor gains.

Within the range of stable feedback, large gain and low damping lead to oscillations, whereas low gain and large damping result in slow compensation for perturbations (as illustrated in the Methods). To determine how sensorimotor gains are affected by an increase in stiffness or neural delay, we first determine the best combination of feedback gains and the resulting balance performance, for a given stiffness and delay.

### 2. Critical feedback gains

We assume that the best combination of feedback gains results in fast compensation of perturbations without oscillations. Subjects typically adopt such feedback gains when responding to lateral perturbations of stance (23). In second order systems governed by a characteristic polynomial 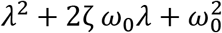, for a given value of *ω*_0_, the fastest compensation without oscillations occurs for the critical damping *ζ* = 1. For such critical damping, the characteristic equation has a unique double root-*ω*_0_. If a perturbation brings the system away from its equilibrium position, then the system is returned to its initial position following the time-course 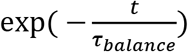 where 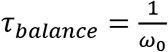. Higher damping results in slower compensation for perturbations, whereas lower damping results in oscillations. The characteristic equation of the system with delayed neural feedback is:

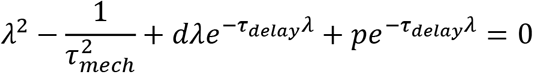

To determine the feedback gains which provide critical damping, we use a linear approximation to the delay introduced by Pade (24): 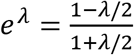. With this approximation, the characteristic equation becomes a third order polynomial, provided in the Methods. Then, we generalized the notion of critical damping to third order systems, considering that a system is critically damped when it has a unique triple negative root *λ*_0_. The characteristic time to return to equilibrium is then 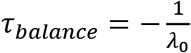. For each value of stiffness and delay, this procedure provided a unique value of *τ_balance_* (Fig 3.A-C), and of the corresponding critical proportional gain *p_crit_* (Fig 3.D-F) and derivative gain *d_crit_* (Fig 3.G-I).

**Fig 3.**
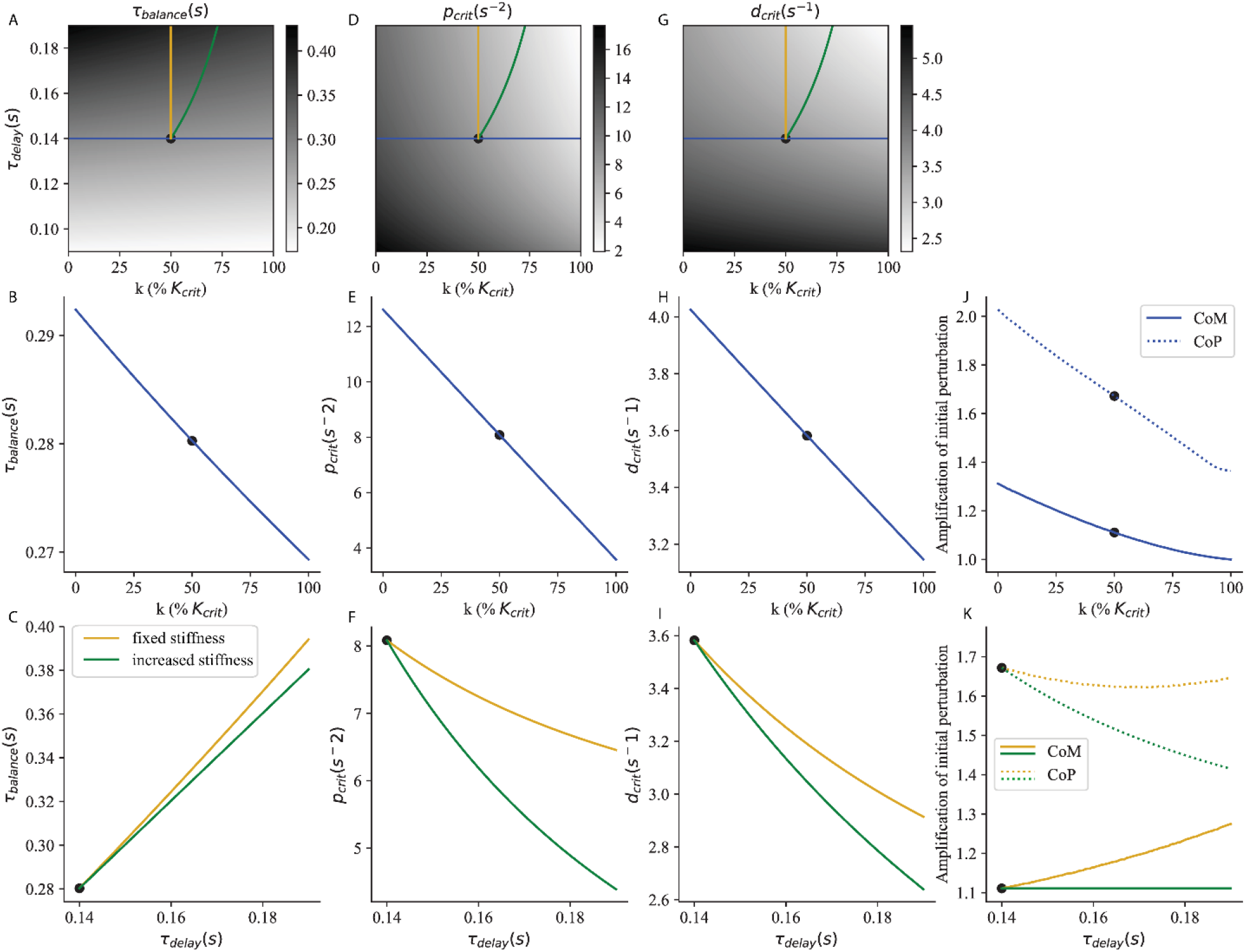
Critical feedback. The time to recover balance (A, B, C), the corresponding critical proportional (D, E, F) and derivative (G, H, I) gains and the peak excursions (J,K) of the CoM (full line) and CoP (dashed line) are shown as a function of stiffness and neural delay. The strategy of increasing ankle stiffness with constant τ_delay_ = 0.14 s is illustrated in blue. The effect of an increase in neural delay with fixed ankle stiffness k = 50% K_crit_ is illustrated in yellow. The strategy of increasing ankle stiffness to compensate for an increased delay (maintaining a constant S) is illustrated in green. The reference system used in section V. is indicated as a black dot.

In Fig 3.A, we show *τ_balance_* as a function of ankle stiffness and neural delay. This corresponds to the time required to recover balance after a perturbation. Light shades of grey correspond to small values of *τ_balance_* and therefore to good balance performance. Balance performance is best for large ankle stiffness and small neural delay (Fig 3.A, lower right corner). For a constant delay, *τ_balance_* decreases with increasing ankle stiffness (Fig 3.A, B, blue curve). For constant ankle stiffness, *τ_balance_* increases with increasing neural delay (Fig 3.A, C, yellow curve). When ankle stiffness and neural delay are jointly increased such that *S* remains constant, this mitigates the increase in *τ_balance_* (Fig 3.A, C, green curve).

The balance performance indicated by *τ_balance_* is only achieved if the sensorimotor gains provide critical damping. The critical proportional (Fig 3.D) and derivative (Fig 3.G) gains are largest for low ankle stiffness and neural delay (lower left corner). For a constant delay, the critical gains decrease with increasing ankle stiffness (Fig 3.D, E, G, H, blue curve). For constant ankle stiffness, the critical gains decrease with neural delay (Fig 3.D, F, G, I, yellow curve). When ankle stiffness and neural delay are jointly increased such that *S* remains constant, this amplifies the decrease in critical gains (Fig 3. D, F, G, I, green curve).

In Figure 3.J, K we show the peak excursion of the centre of mass (CoM, full lines) and centre of pressure (CoP, dashed lines) after a perturbation of arbitrary amplitude 1. For a constant delay, the peak excursion of both the CoM and CoP decrease with increasing ankle stiffness (Figure 3.J). For constant ankle stiffness, with increasing delay the CoM excursions increase (Figure 3.K, full yellow line), whereas the CoP excursions slightly decrease (Figure 3.K, dashed yellow line). When ankle stiffness and neural delay are jointly increased such that *S* remains constant, the CoM excursions remain constant (Figure 3.K, full green line), and the CoP excursions actually decrease (Figure 3.K, dashed green line).

We illustrate the importance of co-adjusting ankle stiffness and sensorimotor gains by simulating how systems with various stiffness sensorimotor gains and neural delays respond to a perturbation.

### 3. Improving stability in challenging balance conditions

In Fig ***4***, we show the response of the system with *τ_delay_* = 0.14 *s* to an external perturbation occurring at time 0, which causes an initial shift in ankle angle by an arbitrary distance 1. We first consider a Reference system without ankle stiffness and with the corresponding critical sensorimotor gains (Fig ***4***.A). The initial perturbation causes an immediate increase in the torque of weight (Fig ***4***.A, black). The ground reaction torque component due to stiffness remains null (Fig ***4***.A, dark blue). Therefore, the position of the centre of pressure matches the ground reaction torque due to contraction (Fig ***4***.A, light blue) and remains immobile during the delay period. The person therefore starts to fall, and picks up speed, as reflected by the increase in the torque of weight (Fig ***4***.A, black) during the delay period. When the feedback control intervenes, a large increase in contraction is therefore necessary to first slow down falling, then return the ankle angle to its initial position.

**Fig 4.**
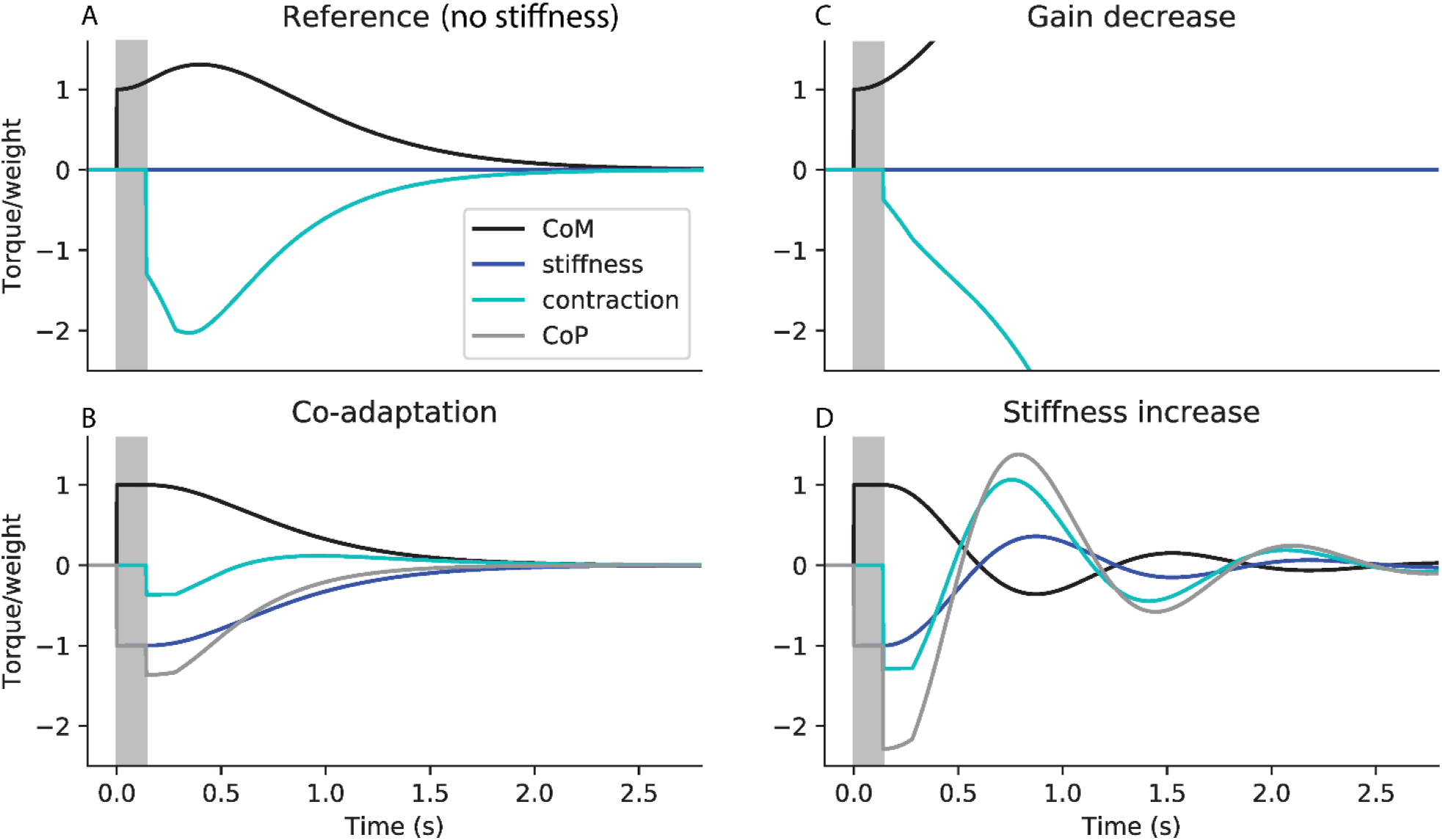
Co-adaptation of ankle stiffness and sensorimotor gains to improve immobility. Simulated response to a perturbation in ankle angle of arbitrary amplitude 1, showing the torque of weight (black, corresponding to the position of the centre of mass), and the ground reaction torque (grey, corresponding to the position of the centre of pressure), with its two components respectively due to stiffness (dark blue) and feedback contraction (light blue). All torques are normalised to weight. A. Reference system without stiffness, and with the corresponding critical feedback. B. Co-adaptation strategy: system with critical stiffness and critical feedback. C. Gain decrease strategy: system without stiffness, and with the same feedback controller as in B. D. Stiffness increase strategy: system with critical stiffness, and with the same feedback controller as in A. The response delay (between the perturbation and the first change in contraction) is shaded in grey. In panels A and B, the CoP is not shown as it overlaps with the torque due to contraction (light blue).

We then consider the co-adaptation strategy, in which stiffness is increased to its critical value, and the sensorimotor gains are decreased to their corresponding critical values (Fig ***4***.B). The initial perturbation causes an immediate increase in the torque of weight (Fig ***4***.B, black). Since ankle stiffness perfectly compensates for the torque of weight, there is an immediate, equivalent and opposite increase in the ground reaction torque component due to stiffness (Fig ***4***.B, dark blue). Therefore, during the delay period (shaded in grey), the CoM remains immobile. When the feedback control intervenes, a small increase in contraction (Fig ***4***.B, light blue) is sufficient to nudge the person back upright. The co-adaptation strategy thus allows perturbations to be cancelled faster, with less overshoot in both the CoM and CoP, and less increase in contraction.

This improvement in performance cannot be attributed to either the decreased gains or the increased stiffness alone. Indeed, if only the gains are decreased, without a corresponding increase in stiffness, then the system is unstable (Fig ***4***.C). The small increase in contraction at the end of the delay period (Fig ***4***.C, light blue) is insufficient to compensate for the large increase in ankle angle and speed (Fig ***4***.D, grey) during the delay period. If, on the other hand, only stiffness is increased without a corresponding decrease in gains, then the system is oscillatory (Fig ***4***.D). During the delay period, the person remains immobile. The large increase in contraction (Fig ***4***.D, light blue) then causes the ankle angle to overshoot its initial position, resulting in oscillations.

### 4. Compensating for increased feedback delays

With ageing, the ankle muscle responses to perturbations of stance are delayed (13,14). Fig ***5*** illustrates the importance of co-adapting ankle stiffness and sensorimotor gains to the neural delay. We consider a Reference system (Fig ***5***.A) with a delay of 0.14 s, intermediate stiffness (k = 50% K_crit_), and the corresponding critical feedback gains. We then consider that the delay is doubled (Fig ***5***. B – D, the additional delay is shaded in red). If the person does not adjust their stiffness (Fig ***5***. B, C), then the CoM (black) falls further and gains more speed during the prolonged delay. If additionally the person does not adjust their feedback gains, then the large CoM excursions in turn cause large CoP excursions (Fig ***5***. B, peak in the grey curve). The CoM therefore overshoots its initial position and the system enters into oscillations.

**Fig 5.**
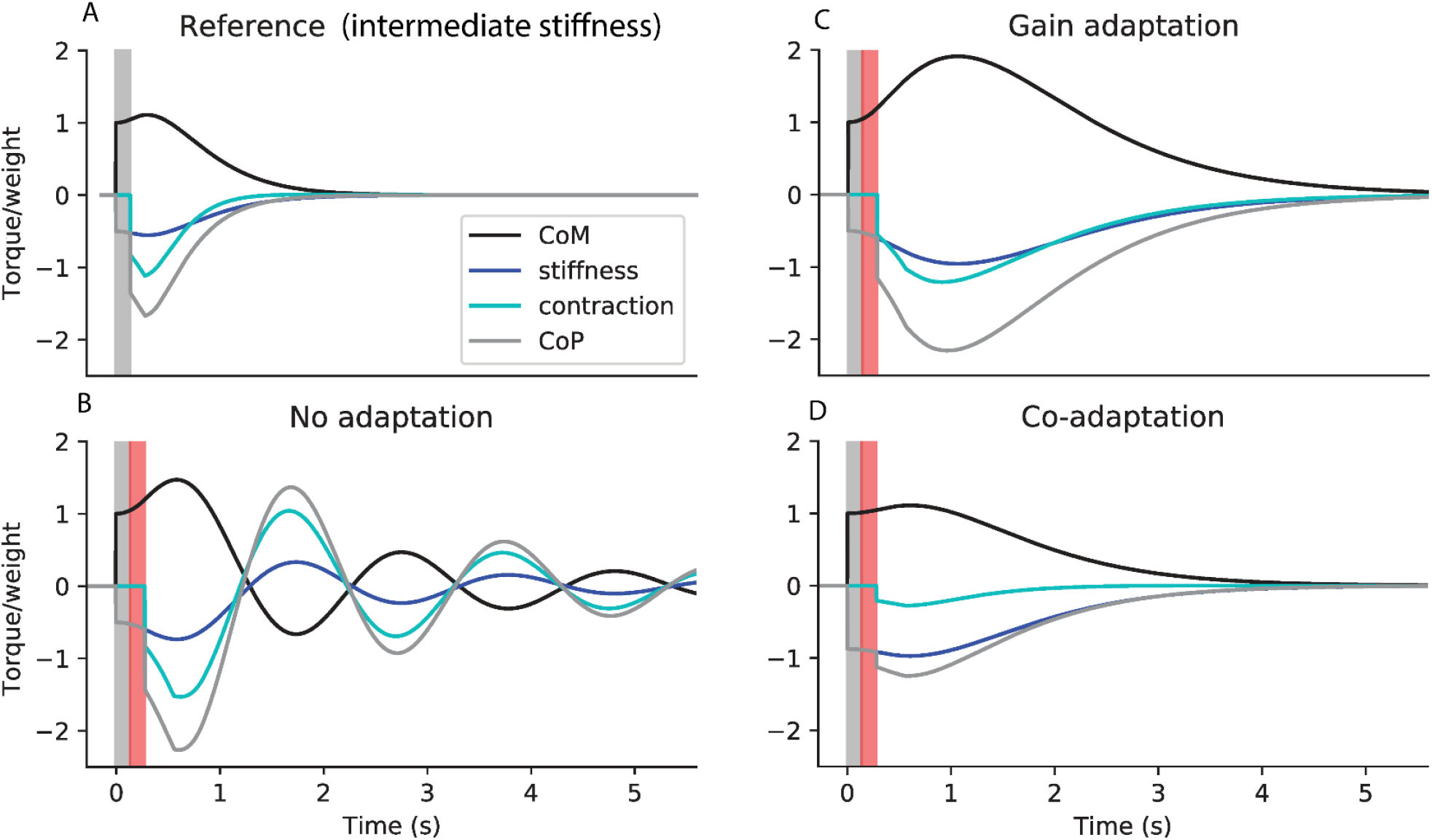
Co-adaptation of ankle stiffness and sensorimotor gains to compensate for increased delays. Simulated response to a perturbation in ankle angle of arbitrary amplitude 1, showing the torque of weight (black), and the ground reaction torque (grey), with its two components of due to stiffness (dark blue) and feedback contraction (light blue). All torques are normalised to weight. A. Reference system with a delay of 0.14 s (shaded in grey in all panels), intermediate stiffness k = 50% K_crit_, and the corresponding critical gains. B.-D. Increase in delay by 0.14 s (shaded in red). B. No adaption: same stiffness and gains as A. C. Gain adaptation strategy: same stiffness as A, and critical gains corresponding to the increased delay. D. Co-adaptation strategy: increase in stiffness to maintain a constant relative speed S, and corresponding critical gains.

Thus, compensating for an increase in neural delay requires a decrease in sensorimotor gains. We illustrate the gain adaptation strategy (Fig ***5***. C), in which the gains are decreased to the critical values appropriate for the increase in delay (Fig ***3***. F, I yellow curve). This prevents the oscillations of the CoM. Relative to the Reference (Fig ***5***.A), both CoM (black) and CoP (grey) excursions are nevertheless increased.

Preventing this increase in CoM and CoP excursions requires a decrease in ankle stiffness. We illustrate the co-adaptation strategy (Fig ***5***. D), in which stiffness is increased such that, at the end of the prolonged delay period, the CoM (black) has fallen just as far as for the Reference system (Fig ***5***. A, black), at the end of its shorter delay period. The gains must be decreased to compensate for the increase in both neural delay and stiffness (Fig 3. F, I green curve). This prevents the increase in CoM excursions with increasing delay (the peak excursion of the CoM in black is identical in Fig ***5***.A and D), and the peak CoP excursion is actually decreased (grey, Fig ***5***. A and D). When the co-adaptation strategy is employed, the main effect of the increase in delay is to increase the time required to cancel the perturbation.

### 5. Generalisation to more complex neuromechanical models

Our modelling results show that, for a given neural delay, it is advantageous to increase the mechanical time constant to slow down body dynamics. Indeed, this decreases the amplification of perturbations during the response delay. In the sagittal plane, this can be achieved by stiffening the ankle. Moreover, our model predicts that when subjects adopt this strategy, they must additionally decrease their sensorimotor gains to prevent over-compensation. Additionally, our results show that following an increase in feedback delay, sensorimotor gains must be decreased. These results generalise to more complex neuromechanical models.

The intrinsic damping properties of muscles are thought to be crucial for motor stabilization (25). We extend our model to include mechanical damping *d_M_* such that:

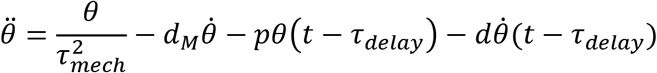

We apply our method for determining critical feedback to this extended model (see Methods), and we find that mechanical damping improves balance performance (decrease in *τ_balance_* in Fig **6**.A, D) and requires a larger proportional gain (Fig **6**.B, E), and a smaller derivative gain (Fig **6**. C, F). Our main result is however not affected: both increased stiffness (Fig **6**.A-C) and neural delay (Fig **6**.D-F) require a decrease in proportional and derivative gains.

**Fig 6.**
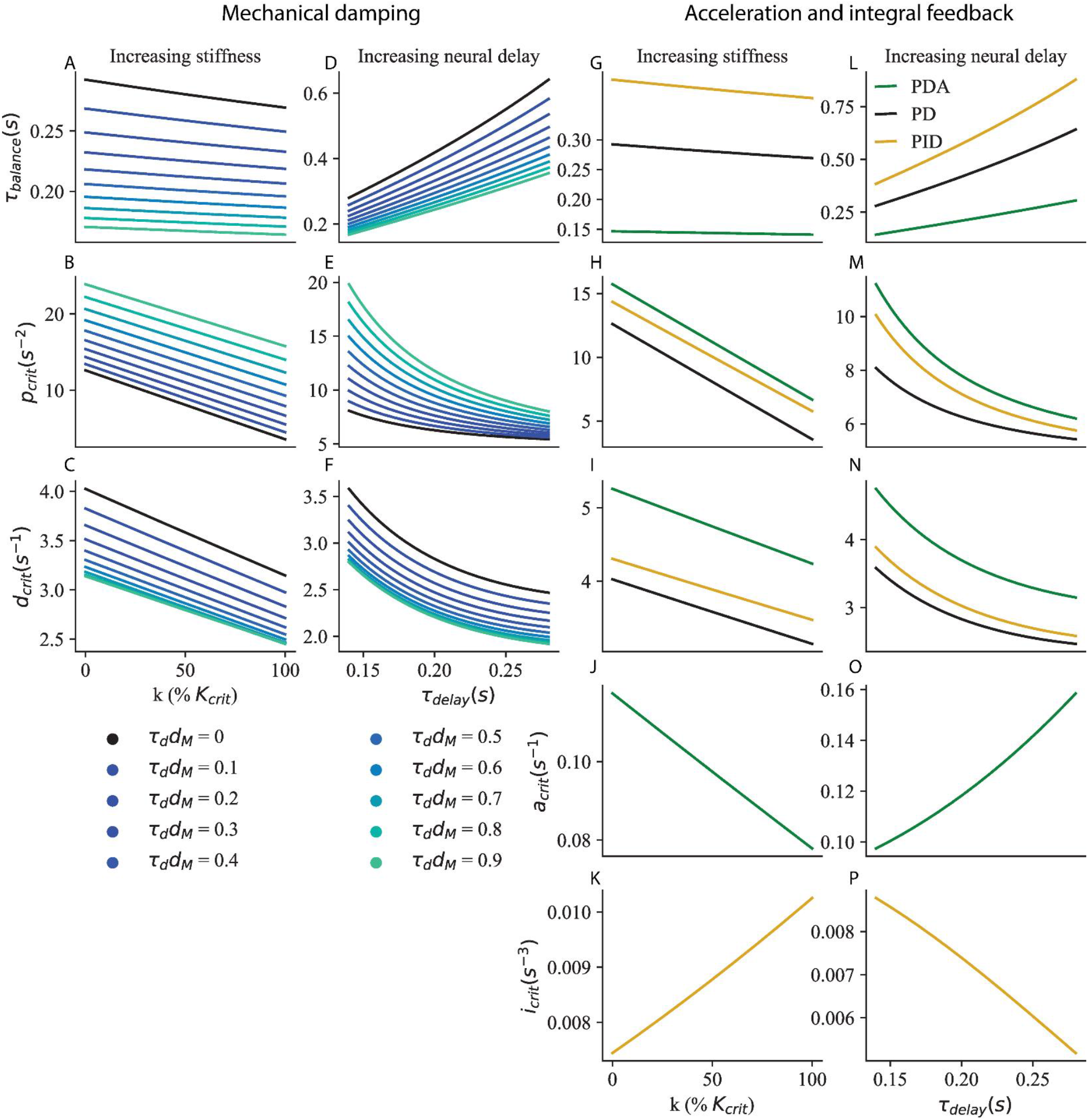
Generalisation to more complex neuromechanical models. A-F Mechanical damping d_M_. The time to recover balance (A, D), critical proportional gain (B, E) and critical derivative gain (C, F) are shown for τ_delay_ = 0.14 s and increasing ankle stiffness (A – C), and for k = 50% K_crit_ ankle stiffness and increasing neural delay (D – F). G-N Acceleration (green) and integral (yellow) feedback. The time to recover balance (G, L), critical proportional gain (H, M), critical derivative gain (I, N), critical acceleration gain (J, O) and critical integral gain (K, P) are shown for τ_deiay_ = 0.14 s and increasing ankle stiffness (G – K), and for k = 50% K_crit_ ankle stiffness and increasing neural delay (L – P).

To fit the muscular contraction responses of human subjects to forwards and backwards perturbations of stance, Welch and Ting found that it was necessary to include acceleration feedback as well as proportional and derivative gains (20). We extend our model to include acceleration feedback *A* such that:

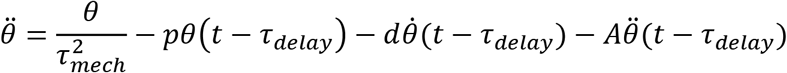

We apply our method for determining critical gains to this extended model (see Methods), and we find that acceleration feedback improves balance performance (decrease in *τ_balance_* in Fig **6**. G, L for PDA control in green relative to PD control in black) and requires a larger proportional gain (Fig **6**. H, M), and derivative gain (Fig **6**. I, N). Our main result is however not affected: both increased stiffness (Fig **6**. G – K) and neural delay (Fig **6**. L – P) require a decrease in proportional and derivative gains. The acceleration gain must be decreased for increasing ankle stiffness (Fig **6**. J) and increased for increasing neural delay (Fig **6**. O).

To better fit the centre of mass (CoM) responses of subjects to visual and platform rotation perturbations, Peterka included integral feedback in their model of standing balance (26). We apply our method for determining critical gains to this model (see Methods), and we find that it increases the time required to cancel perturbations (increase in *τ_balance_* in Fig **6**.G, L for PID control in yellow) and requires a larger proportional gain (Fig **6**.H, M), and derivative gain (Fig **6**.I, N). Our main result is however not affected: both increased stiffness (Fig **6**. G – K) and neural delay (Fig **6**. L – P) require a decrease in proportional and derivative gains. The integral gain must be increased for increasing ankle stiffness (Fig **6**.K) and decreased for increasing neural delay (Fig **6**.P).

Finally, standing balance in the sagittal plane is better modelled by considering not only ankle angle but also hip angle (27) and knee angle (28). In the Methods, we show that our results also generalise to multi-dimensional systems.

## III. Comparison to experiments

### 1. Improving stability in challenging balance conditions

When a subject is asked to stand in a situation in which it is particularly important not to move (Fig 1.B), a decrease in the spinal sensorimotor gain is observed (4–6). This spinal feedback gain can be probed experimentally using the H-reflex (29). This paradigm uses electrical stimulation of the calf muscle nerve to directly excite the sensory fibres embedded within the calf muscle (Fig 1, blue arrows), which are sensitive to the stretch and speed of stretch of the calf muscle, and therefore to both ankle angle and speed. In turn these sensory fibres directly excite the motor neurons of the ankle muscles, located in the spinal cord, which increase the contraction of the calf muscle (Fig 1, red arrow). The change in muscle contraction for a given electrical stimulation is called the H-reflex, and is reduced by around 19% when standing facing the edge of an elevated platform (4), 20% when standing on a narrow platform (5), and 24% when standing with the eyes closed (6). This decrease in spinal gains is usually taken to indicate an increased reliance on the supra-spinal control of balance.

However, our modelling results show that a decrease in spinal gains alone is not beneficial for supra-spinal control. Cortical and other supra-spinal contributions to balance have longer delays than the spinal stretch reflex. During this supra-spinal delay, the dynamics are determined only by the body mechanics and the spinal control. A decrease in spinal gains (without a corresponding increase in stiffness) causes the CoM to fall faster and further during this delay period (as schematically exaggerated in Fig ***4***.B). The effect is therefore to worsen the resulting initial conditions which the supra-spinal control will have to deal with. For example, the effect of a 15% decrease in sensorimotor gain on balance performance is reported in Table 1 (Gain decrease) in absolute values and relative to a Reference system (with a 0.14 s delay, intermediate stiffness 50% K_crit_ and corresponding critical gains, indicated as black dots in Fig 3). The system remains stable, and the decrease in gains enables a 7.9% decrease in CoP excursions. However, the CoM excursions increase by 2.0%. This strategy therefore does not seem appropriate when standing in challenging balance conditions. A cortical control strategy would therefore not benefit from a decrease in spinal gains.

**Table 1.** Standing in challenging balance conditions. For each of the Reference, Gain decrease and Co-adaptation strategies, the Table shows the values of stiffness, neural delay and sensorimotor gains used for the simulation, as well as the peak CoM and CoP excursion after a perturbation of arbitrary amplitude 1. All changes are shown as a percent of the reference value, except stiffness changes, which are reported in % K_crit.

| | Stiffness<br>(% $K_{crit}$) | Delay<br>(s) | p<br>(N.m/(s<sup>2</sup>.rad)) | d<br>(N.m/(s.rad)) | Peak<br>CoM | Peak CoP |
| --- | --- | --- | --- | --- | --- | --- |
| Reference | 50 | 0.14 | 8.08 | 3.58 | 1.11 | 1.67 |
| Gain decrease | 50 | 0.14 | 6.87 | 3.46 | 1.13 | 1.54 |
| Co-adaptation | 63.4 | 0.14 | 6.87 | 3.46 | 1.07 | 1.58 |
| Change relative to the Reference |  |  |  |  |  |  |
| Gain decrease | 0 | 0 | - 15% | - 3.3% | + 2.0% | - 7.9% |
| Co-adaptation | + 13.4%<br>$K_{crit}$ | 0 | - 15% | - 3.3% | - 3.6% | - 5.3% |

There is evidence that subjects indeed increase ankle stiffness when standing in challenging balance conditions by co-contracting ankle muscles. Ankle muscle contraction may thus increase by up to 10% when standing on a narrow platform (5) or when closing the eyes (6), and the contraction of the shin muscle tibialis anterior is increased by up to 50% when standing on an elevated platform (19). Direct measurements of ankle stiffness in such standing conditions are unfortunately not available. When subjects are asked to lean forwards relative to normal stance, the contraction of the calf muscle gastrocnemius increases up to three-fold, with an increase in ankle stiffness by around 15% *K_crit_* (29). The moderate increases in co-contraction observed in challenging balance conditions may thus partially co-adapt ankle stiffness. It is however unclear whether they achieve the 13.5% K_crit_ increase in ankle stiffness required by the co-adaptation strategy.

We suggest an alternative interpretation for the observed decrease in spinal gains. According to our modelling results, the appropriate strategy to decrease CoM and CoP excursions is to simultaneously increase ankle stiffness and decrease gains (Figure 3.B, E, H, J). A 15% decrease in spinal gain would thus improve balance if it were accompanied by an increase in stiffness of 13.5% K_crit_. This co-adaptation strategy enables a decrease in both the CoM and CoP excursions, by respectively 3.6% and 5.3% (Table 1, Co-adaptation). It is thus an appropriate strategy for improving immobility in challenging balance conditions.

The human body comprises joints other than the ankle which are relevant for balance, such as the hip and knee joints (28). Our modelling results generalise to multi-dimensional systems and therefore predict that standing balance may also benefit from the stiffness of muscles acting at joints other than the ankle, if they slow down the amplification of perturbations during the response delay. This has indeed been shown by De Groote and colleagues, who asked human subjects to stand still despite external perturbations (30). They observed the resulting motion of the body during the time it takes for the nervous system to intervene, and attempted to reproduce this motion in simulations. They found that if muscle stiffness was not included in their simulations, then the simulated body fell much faster than the human subjects. Thus, human leg muscles are arranged in such a way that muscle stiffness slows down falling. This suggests that further increases in the stiffness of leg muscles (through co-contraction) may be a useful strategy to further decrease falling speed. This would then require a decrease in sensorimotor gains.

### 2. Compensation for increased feedback delays

With aging, there is an increase in the H-reflex latency of the ankle muscle soleus by 10 to 15% (10,11). The response of ankle muscles to support surface rotations and translations is delayed by up to 30% (13,14). We consider the effect of an intermediate 20% increase in delay on balance performance. If the subject does not change their sensorimotor gains (No adaptation), due to the increase in delay, the movement of the CoM becomes oscillatory (schematically exaggerated in Fig 5.B). Additionally, the peak excursions of both the CoM and CoP are increased, respectively by 4.4% and 5.7% (Table 2, No adaptation). To prevent the CoM oscillations, subjects must decrease their gains to the new critical values (Table 2, Gain adaptation). This also has the effect of decreasing the CoP excursions by 2.9% relative to the Reference. However, CoM excursions are increased by 7.1%. Maintaining the same CoM excursions as the Reference requires a combined increase in ankle stiffness and decrease in gains (Co-adaptation). This additionally provides a decrease in peak CoP excursions by 10.3% (Table 2, Co-adaptation).

**Table 2.** Compensating for increased delay. The Table reports the sensorimotor control parameters and performance of the Reference, and the three strategies for increased delay (No adaptation, Gain adaptation and Co-adaptation). All changes are shown as a percent of the reference value, except stiffness changes, which are reported in % K_crit.

| | Stiffness<br>(% $K_{crit}$) | Delay<br>(s) | p<br>(N.m/(s<sup>2</sup>.rad)) | d<br>(N.m/(s.rad)) | Peak<br>CoM | Peak CoP |
| --- | --- | --- | --- | --- | --- | --- |
| Reference | 50 | 0.14 | 8.08 | 3.58 | 1.11 | 1.67 |
| No adaptation | 50 | 0.168 | 8.08 | 3.58 | 1.16 | 1.76 |
| Gain adaptation | 50 | 0.168 | 6.99 | 3.15 | 1.19 | 1.62 |
| Co-adaptation | 65.3 | 0.168 | 5.61 | 2.99 | 1.11 | 1.50 |
| Change relative to the Reference |  |  |  |  |  |  |
| No adaptation | 0 | + 20% | 0 | 0 | + 4.4% | + 5.7% |
| Gain adaptation | 0 | + 20% | - 13.5% | - 12.2% | + 7.1 % | - 2.9% |
| Co-adaptation | + 15.3%<br>$K_{crit}$ | + 20% | - 30.6% | - 16.7% | 0 | - 10.3% |

The co-adaptation strategy requires a 30.6% decrease in proportional gain. This is consistent with the 30 to 36% decrease in soleus H-reflex occurring during aging (7–9). The muscular contraction response to platform translations is likewise strongly reduced (14). The co-adaptation strategy also requires an increase of stiffness by 15.3% K_crit_. The ankle stiffness measured with relaxed muscles may be more than 20% (31) (more than 10% for a single foot). Additionally, co-contraction during quiet standing is approximately doubled in older subjects (7,12). To the best of our knowledge, the effect of co-contraction on ankle stiffness has not been quantified in older subjects. However it should further increase ankle stiffness relative to relaxed muscles. It is therefore plausible that the 15.3% K_crit_ increase in ankle stiffness required by the co-adaptation strategy is achieved during normal ageing.

The observed changes in spinal delay, gain and ankle stiffness with aging are thus consistent with the co-adaptation strategy. The decrease in spinal gain with aging therefore does not necessarily reflect an increased cortical control of balance (34), but rather an adaptation for increased delays.

## IV. Discussion

### 1. Anticipatory co-adjustment of mechanical properties and sensorimotor gains for motor stabilisation

We have presented a generic model of stabilisation with neural response delays, which highlights the importance of co-adapting ankle stiffness and sensorimotor gains. In situations in which it is particularly important to remain immobile (such as standing on a narrow support, or in front of a cliff), our model suggests that the most effective way to improve balance is to increase ankle stiffness and decrease sensorimotor gains. Indeed, the increase in ankle stiffness slows down the CoM fall during the neural response delay. As the CoM falls more slowly, a smaller increase in contraction is then required to return it to its initial position. This co-adaptation strategy allows perturbations to be cancelled faster, and with smaller excursions in both the CoM and CoP. We have reviewed experimental evidence that, in challenging balance conditions, young subjects indeed decrease spinal gains and at least partially increase ankle stiffness.

During aging, the neural response delay to perturbations of stance increases. According to our modelling results, an increase in delay without adaptation of either stiffness or gains results in oscillations of the CoM, as well as increased excursions of both the CoM and CoP. To prevent such oscillations, the sensorimotor gains must be decreased. This gain adaptation strategy prevents CoM oscillations, and decreases CoP excursions, but further increases the CoM excursions. According to our model, the most effective way to maintain balance performance with increasing neural delay is to increase stiffness and decrease gains. This co-adaptation strategy prevents the increase in CoM excursions, and further decreases the CoP excursions. We have reviewed experimental evidence that in quiet standing, the spinal gains of older subjects are decreased relative to young subjects, and their ankle stiffness is increased.

Interestingly, the co-adaptation strategy not only reduces the CoM excursions (or prevents their amplification with increasing delays), it also reduces the CoP excursions. In the simulations we have presented, the CoP range is assumed to be unlimited. In human stance however, the position of CoP is limited to the extent of the foot. By reducing the CoP excursions after a perturbation of a given amplitude, the co-adaptation strategy allows larger perturbations to be compensated for without requiring unrealistic torques. This may be particularly important for older subjects who have a reduced range of CoP motion (32,33). In practice, for large perturbations requiring unrealistic torques, subjects may complement the spinal balance strategy by taking a step (34). In any case, our model is inappropriate to account for the responses of subjects to large perturbations. Indeed, our analysis is based on a linearization of the dynamics around the equilibrium position. We thus considered that ankle stiffness is constant, however for larger perturbations the ankle stiffness may vary (35).

### 2. Relation to supra-spinal control

The decrease in spinal gains observed when standing in challenging balance conditions, or during aging, is commonly considered as the sign of an increased cortical control of balance. However, our modelling results show that decreasing spinal gains (without co-adapting ankle stiffness) does not provide any advantage for the cortical control of stance in challenging balance conditions. As supra-spinal control has longer delays than spinal control, if the spinal gain is decreased, then the CoM falls further and picks up more speed by the time the supra-spinal control intervenes. This increases the difficulty of the task for the supra-spinal control. The co-adaptation strategy on the contrary reduces the CoM excursions. Thus, independently of supra-spinal control, the task of standing balance becomes easier if spinal gains are decreased in combination with an increase in stiffness. We have also shown that a decrease in spinal gains is an effective strategy to compensate for age-related increases in delays, independently of any assumptions about supraspinal control. According to our model, the most effective strategy is again a combined increase in stiffness and decrease in gains. We therefore argue that the reduction in stretch reflex observed during aging and in challenging balance conditions is not the hallmark of increased cortical control, but rather reflects an increased reliance on ankle stiffness.

Brainstem- or cortically-mediated visual and vestibular responses to perturbations of stance also play an important role for maintaining balance (26,36). These can complement spinal control, particularly when the sensory signal of ankle angle is poorly informative of the CoM state, such as when standing on a platform which rotates the toes up and down (26). We expect our modelling results to apply to visual and vestibular contributions to balance. We therefore predict that, in conditions leading subjects to adopt larger ankle stiffness, visual and vestibular gains are decreased (36). We also predict that visual and vestibular gains are decreased for increased latency.

### 3. Relation to previous models of balance

Previous modelling studies have determined the stability regions for the feedback gains of the delayed PD controller (see equations in the Methods). In a model of lateral stance, Bingham and colleagues have analysed how these regions change as a function of stance width (23). They showed that, for increasing stance width, the region of stable gains shifts towards smaller values, and does not fully overlap with the region of stable gains for a narrow stance width. Similarly, we find that the regions of stable gains do not fully overlap for changing ankle stiffness and neural delay.

Subjects may in principle adopt any of the gains within the stable region, however in practice they adopt gains which lead to a fast compensation of perturbations without overshoot (20). We propose a novel method to determine which combination of gains provides this behavior, and we refer to this combination as the critical gains. We used this method do determine how sensorimotor gains should adjust to changes in stiffness and delay.

A variety of models have been proposed to account for standing balance. Models which focus on the contribution of body mechanics to stance typically include ankle stiffness and may also include damping (37). Different variations may include additional joints, such as the hip and knee joints (27,28). Models which emphasize the sensorimotor contributions typically include delays, as well as the proportional and derivative gains which are necessary for stability (21). Different variations may include acceleration feedback (20) or integral feedback (26).

We have applied our method for determining critical gains to each of these variations. Our main modelling result is robust to these model variations: the most effective strategy to improve balance performance, or to maintain balance performance despite increasing delays, is the co-adaptation strategy, with a combined increase in ankle stiffness and decrease in the proportional and derivative gains.

This demonstrates the importance of including ankle stiffness in models of standing balance to account for experimentally observed balance strategies. Thus, Maurer and Peterka fit a model of standing balance without ankle stiffness (and with integral feedback) to quiet standing sway measures in young and older subjects (38). They account for the changes in sway with increasing age through an increase in the proportional gain (+ 18%), a decrease in feedback delay (−2.9%) and a large increase in the noise injected into the model (+ 50%). The estimated increase in gain and decrease in delay are however incompatible with the observed age-related decrease in amplitude and increase in delay of the responses to perturbations of stance. In a later paper, they suggest that if a subject relies on large ankle stiffness, then the model fit will produce an artificially large value for the gain and an artificially low value for the delay (39). Our modelling results suggest that older subjects rely more on ankle stiffness in quiet standing, and this may explain the puzzling age-related changes in their model fits (38).

### 4. Do subjects maintain critical gains?

The critical sensorimotor gains are close to the minimal gains required for stability (Fig 8.B, C). If a subject is able to precisely maintain the critical gains, then they can cancel perturbations with minimal excursions in the CoM. However, if they are unable to maintain such critical gains due to uncertainty in the process parameters (such as inertia, stiffness or delay) or sensory or motor error, then they may inadvertently adopt gains which are too low for stability. When standing at a large height, this may have catastrophic consequences. Therefore, at very high postural threat, perhaps subjects prefer to adopt robust gains near the middle of the region of stable gains (Fig 8. A), rather than critical gains which minimise CoM excursions.

This might explain the recent experimental observations that spinal gains are actually increased rather than decreased when standing at a very large height. When subjects stand at the edge of an elevated platform of 1.6 m height, the H-reflex is decreased (4). The longer latency (120 – 220 ms) response to a rotation of the ankle joint is however increased (40). Moreover, when the subject stands at the edge of a 3.2 m high platform, the increase in H-reflex is no longer observed and the muscular contraction response to a mechanical tap of the Achilles tendon (called T-reflex) is actually increased (41), suggesting an increase in spinal gains. A potential confound is that, in the second study, subjects wore ankle braces which may alter their use of ankle control for balance. However, in a further study, the T-reflex was also found to be increased on a 3.5 m high platform when the subjects did not wear ankle braces (42).

When standing normally, it might also be the case that young subjects adopt robust gains which are larger than the critical gains. In this case, the decrease in H-reflex observed in challenging balance conditions may reflect the subject’s attempt to adopt gains which are closer to the critical gains, thus improving the immobilisation of the CoM.

The relation between H-reflex and muscle contraction has been extensively studied in a different experimental paradigm (43). In this task, the subject is seated and asked to maintain a constant ankle torque using visual feedback. The ankle is then abruptly rotated, which elicits a burst of muscle contraction. When the subject is asked to maintain a larger initial torque, the stretch contraction is larger. In this task, the stretch reflex thus increases with increasing background contraction, a phenomenon known as gain scaling. When standing however, the H-reflex decreases with increasing muscle co-contraction. Although the H-reflex and stretch reflex cannot be directly compared, this suggests that spinal gains are flexibly adjusted in accordance with the task and context.

### 5. Do subjects maintain critical ankle stiffness?

Ankle stiffness during stance is measured by imposing a rotation of the ankle and measuring the immediate change in ground reaction torque that ensues. Experimental measures vary from 40% to 90% of the critical stiffness (22,35,44,45). The observation that ankle stiffness is typically lower than the critical stiffness has given rise to a controversy on the relative importance of “passive” ankle stiffness and “active” neural feedback control for standing balance (37,46). Common to the two conflicting views is the assumption that ankle stiffness is a fixed mechanical property. These approaches thus fail to take into consideration the subjects’ ability to modulate ankle stiffness through ankle muscle co-contraction (18).

When standing in normal conditions, young subjects typically do not co-contract their ankle muscles; however, they are able to do so when it is particularly important to remain immobile (5,6,19). Likewise, during quiet standing, subjects continuously shift their position slightly over a range of around a centimetre; if however they are explicitly instructed to remain as still as possible, then this range is divided by two (47). This suggests that in normal standing conditions, the standing posture is not adjusted to maximise stability. Specifically, young subjects seem to adopt a value of ankle stiffness which is lower than the critical value, as indicated by their ability to reduce sway by half when instructed to do so (47). One reason to maintain low ankle stiffness is that high ankle stiffness may hinder mobility, as it causes the ground reaction torque to immediately and mechanically cancel the torque of weight. Initiating a movement however requires a net external torque to accelerate the movement (48). We have indeed recently developed a theory of postural control (48), according to which, during normal stance, posture is adjusted in view of mobility rather than immobility. Thus, during normal standing in young adults, stability is not maintained at its maximal possible value, possibly due to trade-offs with both mobility and fatigue. However, young subjects can and do transiently increase their stability when this becomes important for motor performance, by adjusting the mechanical properties of their body in anticipation of the task.

When measured with relaxed muscles, ankle stiffness is larger in older adults relative to young subjects (31). Additionally, older subjects co-contract ankle muscles even during normal stance (7,12). This suggests a stronger emphasis on stability in older subjects than in young subjects. We suggest that this is a functional compensation for age-related increases in neural delays. However, since older subjects use co-contraction already for normal stance, they may be less able than young adults to transiently increase ankle stiffness in challenging balance conditions. Their overall balance may thus be affected. There is indeed an increasing incidence of falls with ageing (49). Moreover, the increase in stiffness of the passive tissues around the ankle joint suggests that older subjects may be less able than young subjects of transiently decreasing ankle stiffness at movement initiation. Their overall mobility may thus be affected. There is indeed an increasing incidence of mobility impairments with aging (50). Thus, although the age-related increase in ankle stiffness may allow older subjects to preserve stability in normal standing conditions, it may not be sufficient to ensure stability in challenging balance conditions, and it may come at the cost of a decrease in mobility.

## V. Methods

### 1. Mechanical model of stance: the single inverted pendulum

We consider the widely used model of human stance in the sagittal plane: the single inverted pendulum model presented in **Fig 7** (20,21). When someone is standing on the ground, there are two external forces exerted on them: their weight and the ground reaction force (**Fig 7**). The point of application of the person’s weight is called the centre of mass, noted CoM. The torque of weight around the person’s ankles is thus *mg*L sin(*Θ*) (with *m* the person’s mass, *g* gravity, L the CoM height and *Θ* the ankle angle). To understand the relative roles of body mechanical properties and feedback control in stabilisation, we decompose the ground reaction torque into a mechanical component *K*(*Θ*) due to ankle stiffness, which changes immediately upon a change in ankle angle, and a component *C* due to feedback muscle contraction with only changes after a delay *τ_delay_* after a mechanical perturbation. The sum of the external torques affects the person’s rotational momentum 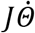, where J is the rotational inertia around the ankles, according to the following equation:

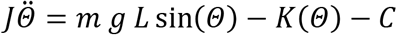

**Fig 7.**
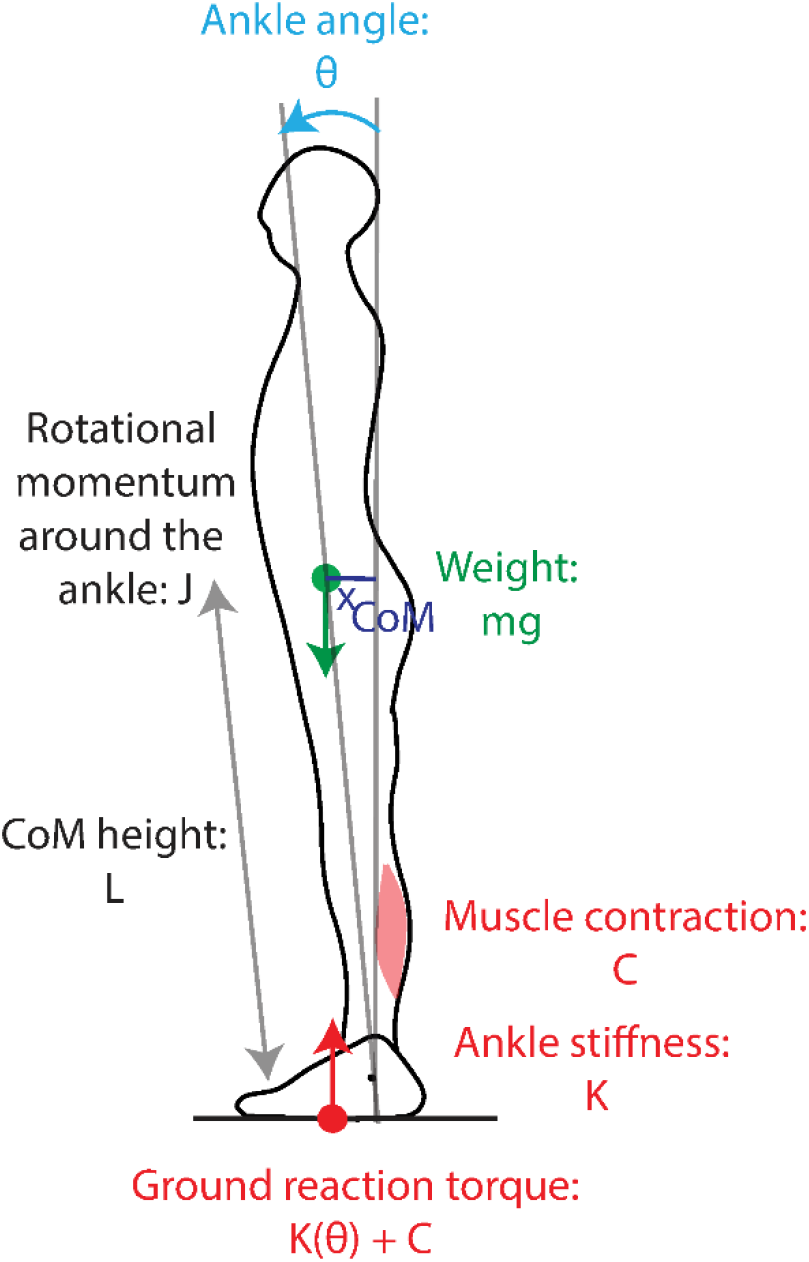
Single inverted pendulum model of stance. The external forces acting on the body are the weight (green arrow), whose torque around the ankle depends on ankle angle θ, and the ground reaction force (red arrow) whose torque depends on both ankle angle and muscle contraction.

We consider that the person is initially at equilibrium at an angle *Θ*_0_ with muscular contraction *C*_0_, and linearize around this equilibrium, introducing *θ, k, c* such that:

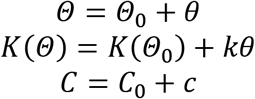

After linearization, the change in rotational momentum becomes:

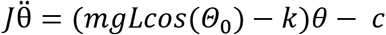

The dynamics thus follows:

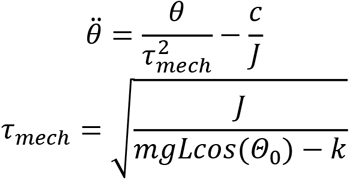

The lower bound of *τ_mech_* corresponds to *k* = 0. With mechanical parameters corresponding to human stance (*J* ≈ *mL*^2^, *L* ≈ 1 *m*, cos(*θ*_0_) ≈ 1), the minimal mechanical time constant is thus:

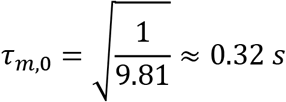

For the neural feedback control, we consider a minimal time delay *τ*_*d*,0_ = 0.14 *s*.

Suppose an initial perturbation shifts the CoM position *θ* and speed 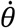 away from their initial equilibrium position to arbitrary values *θ*(*t* = 0) = *θ*_0_, and 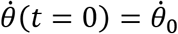. During the neural response delay, the CoM position follows the time course:

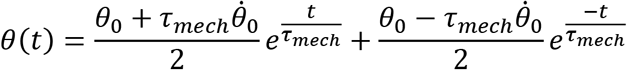

Thus, by the time the nervous system intervenes at *t* = *τ_delay_*, the initial perturbation amplitude *θ*_0_ has been amplified to the value 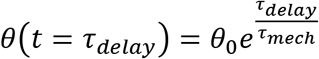, and the initial perturbation speed has been amplified to 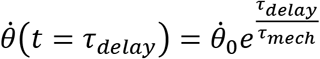.

### 2. Stability analysis

We consider that the neural response follows a delayed proportional-derivative feedback controller, such that the dynamics of the system follows:

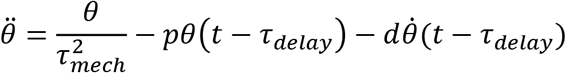

The characteristic equation of the system is therefore:

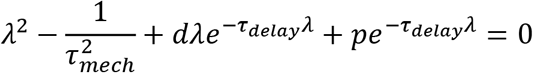

The stability boundaries for this equation have been previously published (21). A derivation based on the Nyquist criterion is provided in S1.I. The boundary consists in two curves (Fig 8.A). The first is the line 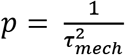. The second is a curve parametrised by *ω* > 0:

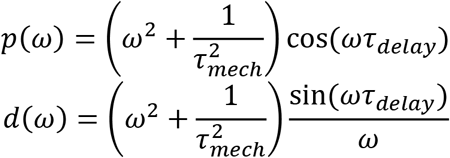

**Fig 8.**
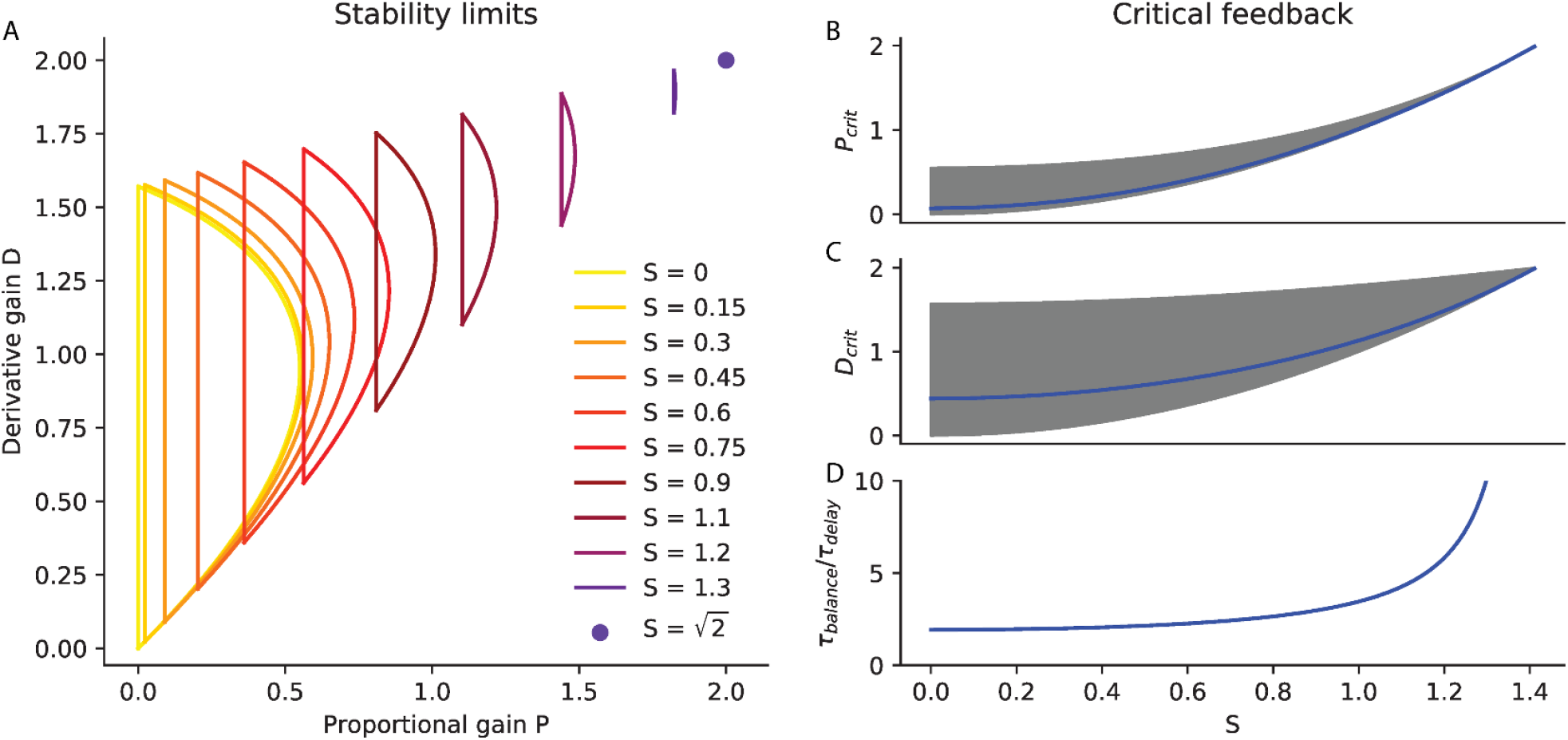
Delayed feedback control. A. Stability limits. The range of stable normalised proportional and derivative gains (P and D) is plotted for different values of relative speed S ranging from S = 0 to the maximal 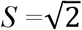. B-D Critical feedback. B. The critical proportional gain P_crit_ is plotted in blue as a function of S. The range of stable proportional gains is shaded in grey. C. The critical derivative gain D_crit_ is plotted in blue as a function of S. The range of stable derivative gains is shaded in grey. D. The time required to recover balance with critical feedback is plotted as a function of S.

We introduce the dimensionless variables: *x* = *ωτ_delay_*, 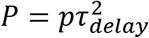, *D* = *dτ_delay_*, 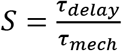. The second curve is then equivalent to the following curve, parametrised by *x* > 0:

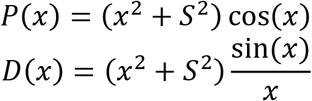

In Fig 8.A, we show the range of feedback gains which can stabilise the system over a range of relative speeds *S*. For a given *S*, the stability limits for the proportional gain *p* scale with 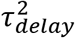 and those for the derivative gain *d* scale with *τ_delay_*. We therefore plot the stability limits in terms of the dimensionless gains 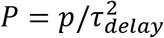 and *D* = *d*/*τ_delay_*. For each value of *S*, there is a limited range of feedback gains (*P, D*) that can stabilize the system. This range is larger for slower values of *S*, suggesting slower systems are more robust to an inappropriate calibration of the feedback gains. Additionally, with increasing *S*, the range of stable gains shifts towards higher values. This shows that if the system speed *S* changes, then the feedback gains must be adjusted to maintain stability.

### 3. Critical feedback gains

The characteristic equation of the linearized inverted pendulum with delayed feedback control is:

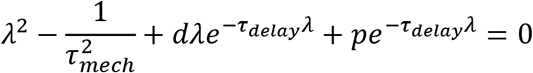

We introduce the dimensionless variable *X* = *τ_delay_λ*. The roots of the characteristic equation are given by:

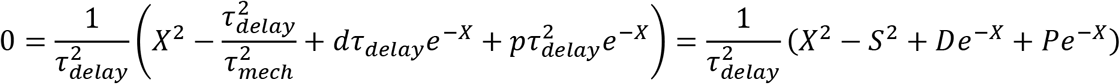

Critical damping is defined for second order systems governed by a characteristic equation of the form:

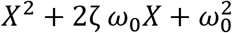

To define critical damping for this system, we first introduce a rational function approximation for the delay, first proposed by Pade (24). With this approximation, the characteristic equation becomes a third order polynomial. We then generalize the notion of critical damping from second order to higher order polynomials.

#### a) Pade approximation

The first order Pade approximation of the delay is given by:

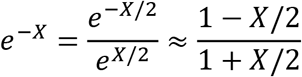

A dynamical systems interpretation of this approximation is provided in S1.II. With this approximation, the characteristic equation becomes:

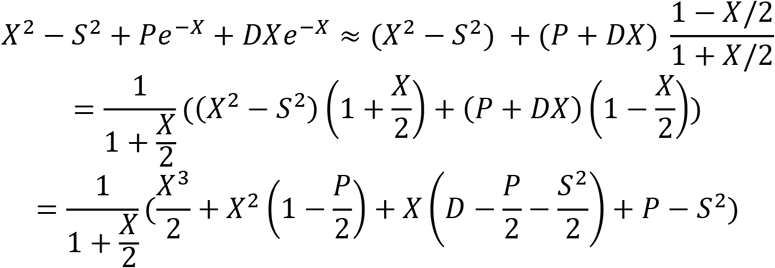

The roots of the approximate characteristic equation are thus the roots of:

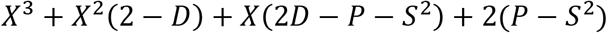

#### b) Generalisation of criticality to higher order polynomials

In second order systems governed by a characteristic equation 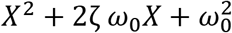, for a given value of *ω*_0_, the fastest compensation without oscillations occurs for the critical damping *ζ* = 1. For such critical damping, the characteristic equation has a unique double root −*ω*_0_. Higher damping results in slower compensation for perturbations, whereas lower damping results in oscillations. We generalize the notion of ‘critical damping’, and consider that critical feedback in an n^th^ order system corresponds to the system having a unique negative root −*ω*_0_ of order n.

#### c) Calculation

We therefore consider that our system is at criticality when is has a unique triple negative root −*ω*_0_. This requires:

The coefficients of the characteristic equation must therefore correspond to the coefficients of the polynomial:

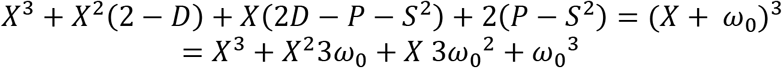

To find the critical feedback gains P_crit_, D_crit_ for a given relative speed S, we solve for (*ω*_0_, P_crit_, D_crit_) the system of equations:

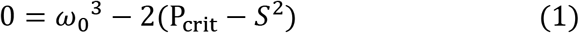

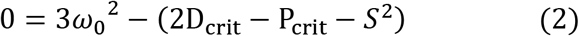

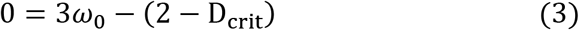

First we determine *ω*_0_ as a function of *S* by removing (*P, D*) from the equations:

According to (3): D_crit_ = 2 – 3*ω*_0_
According to (2): 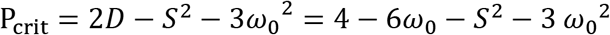
Replacing in (1):

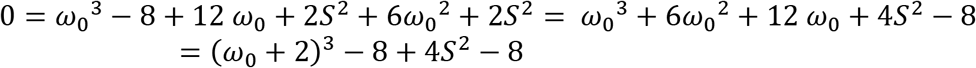
Thus: (*ω*_0_ + 2)^3^ = 4(4 – *S*^2^) > 0

This equation admits one real positive solution for (*ω*_0_ + 2), and two complex conjugate solutions. We take the real solution:

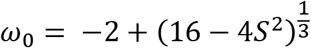

Replacing in (3): 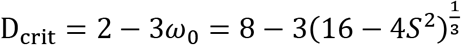
Replacing in (2): 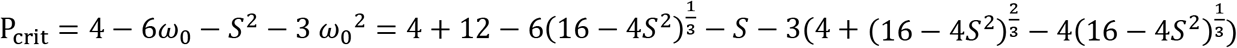

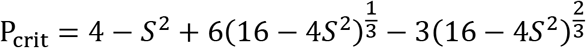

#### d) Illustration

The response to a perturbation of the system with critical derivative gain and various values of the proportional gain is shown for *τ_delay_* = 0.14 *s* with critical stiffness (Fig 9.A) and without stiffness (Fig 9.B); and with a 20 % increase in delay, without stiffness (Fig 9.C). Proportional gains below the critical value (blue) result in slow compensation for perturbations. Proportional gains above the critical value (red) result in oscillations. These results are exacerbated without ankle stiffness and with increasing delay. This may even lead to instability.

**Fig 9.**
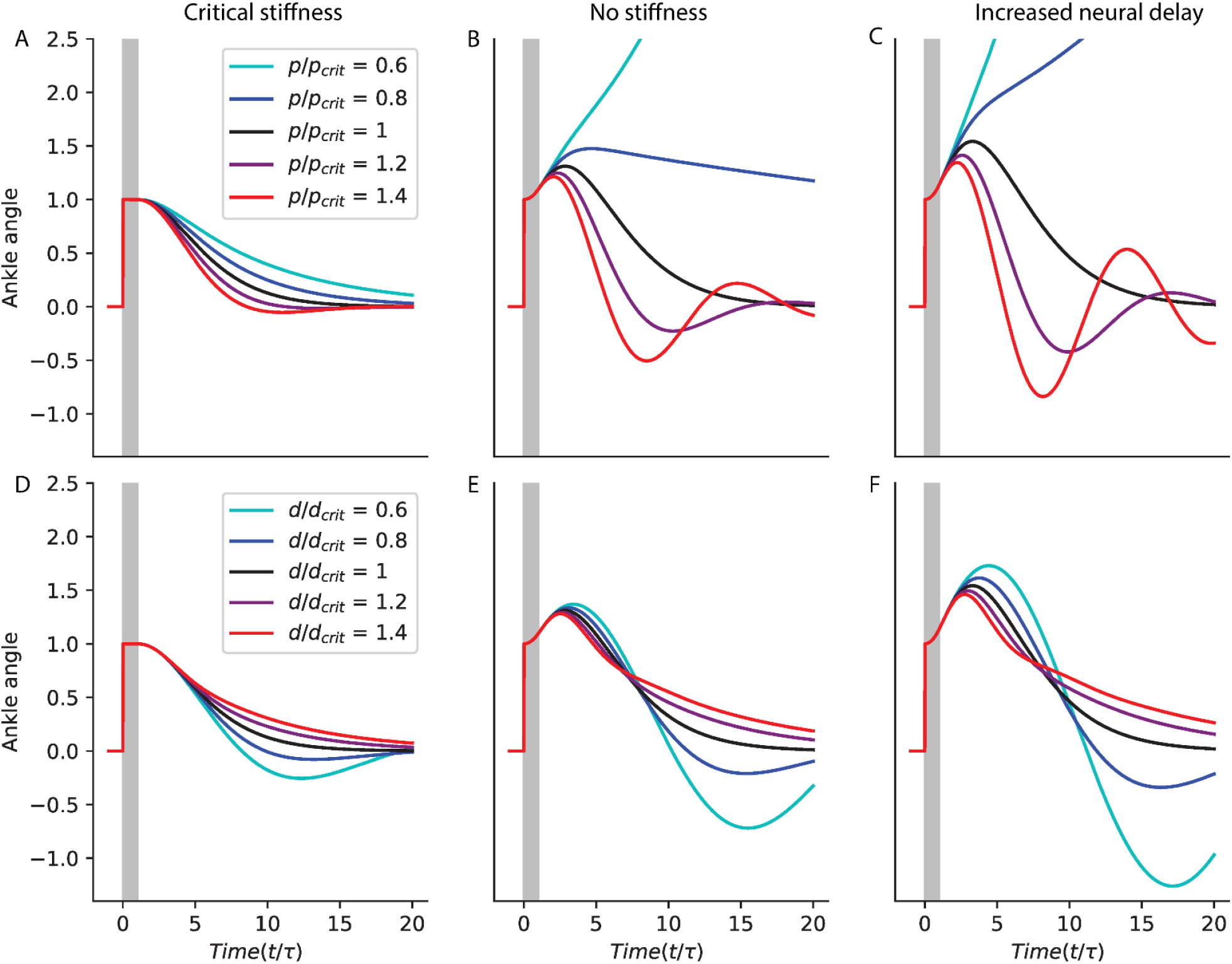
Illustration of critical feedback. Simulated response to a perturbation with critical derivative gain and various values of the proportional gain (A-C); and with critical proportional gain and various values of the derivative gain (D-E); for a system with τ_delay_ = 0.14 s (A, B, D, E) and either critical stiffness (A, D) or no stiffness (B, E); and for a system with a 20% increase in delay and no stiffness (C, F).

The response to a perturbation of the system with critical proportional gain and various values of the derivative gain is shown for *τ_delay_* = 0.14 *s* with critical stiffness (Fig 9.D) and without stiffness (Fig 9.E); and with a 20 % increase in delay, without stiffness (Fig 9.F). Derivative gains below the critical value (blue) result in oscillations. Derivative gains above the critical value (red) result in slow compensation for perturbations. These results are exacerbated without ankle stiffness and with increasing delay.

The critical gains calculated thus provide the fastest compensation of perturbations without overshoot.

#### e) Mechanical damping

We consider that the system has mechanical damping *d_M_* such that:

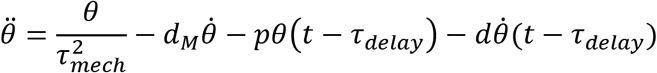

With the dimensionless damping *D_M_* = *τ_delay_d_M_*, the characteristic equation becomes:

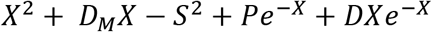

With the Pade approximation, its roots are those of:

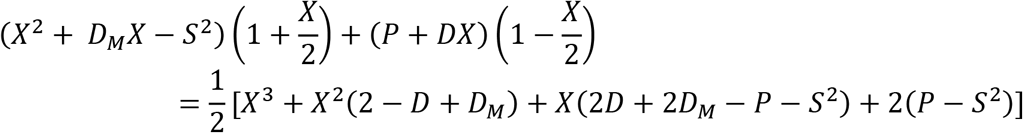

For a given speed *S* and mechanical damping *D_M_*, we numerically solve for (*ω*_0_, P_crit_, D_crit_) the system of equations:

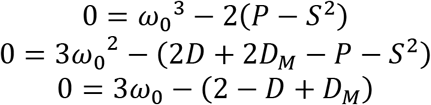

The resulting time to recover balance and critical gains as a function of S are shown in Fig 10.A-C for increasing values of damping.

**Fig 10.**
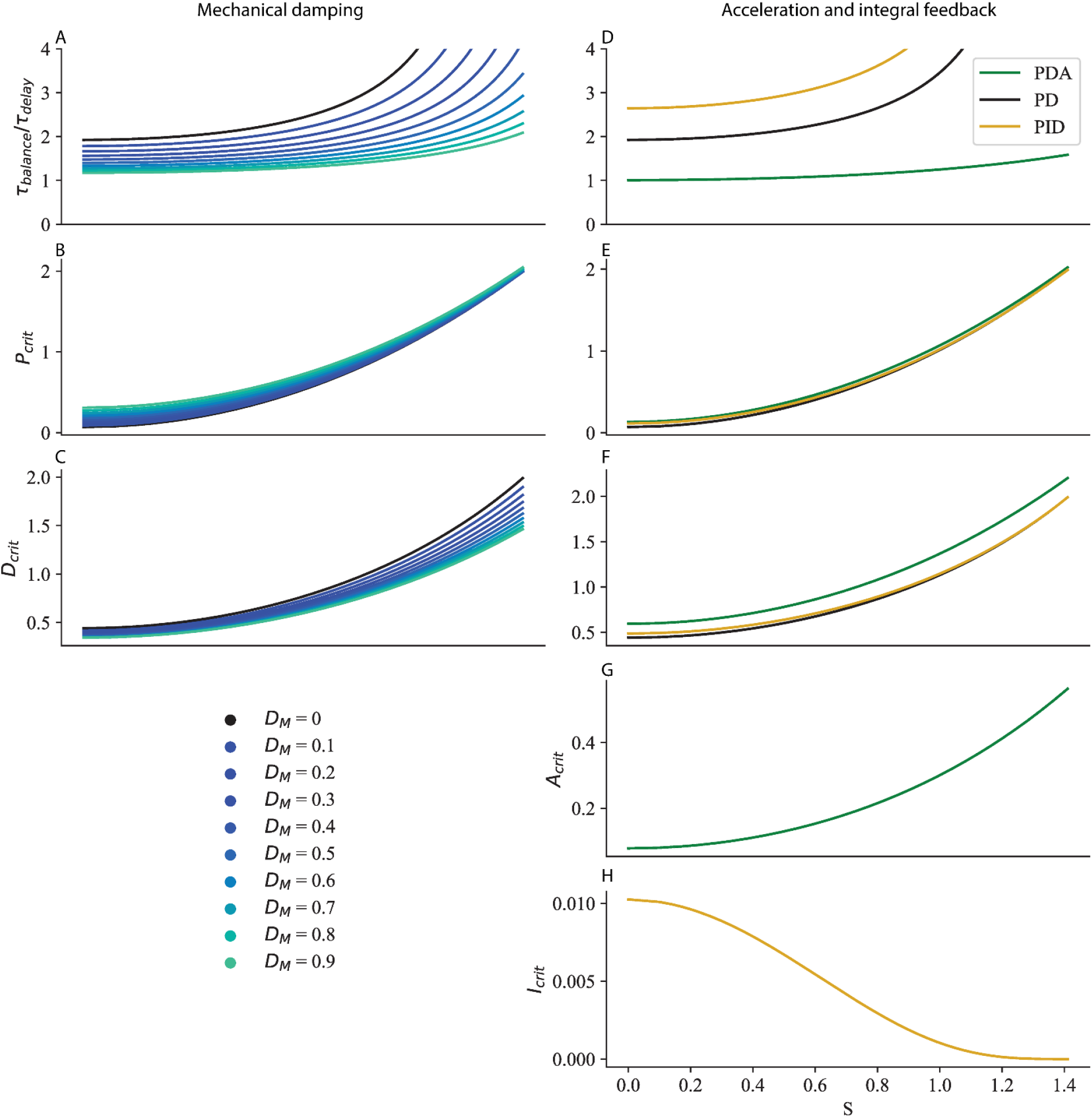
Mechanical damping and acceleration and integral feedback. Time to recover balance (A, D), critical proportional gain (B, E), critical derivative gain (C, F) critical acceleration gain (G) and critical integral gain (H) for increasing values of damping (A-C) and with acceleration (D-G, green) and integral (D, E, F, H, yellow) feedback.

#### f) Acceleration feedback

We consider that the system has acceleration feedback *A* such that:

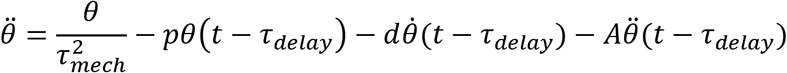

To calculate the critical feedback gains with acceleration feedback *A*, we must use the second order Pade approximation:

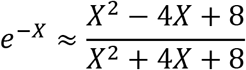

With this approximation, the roots of the characteristic equation are given by:

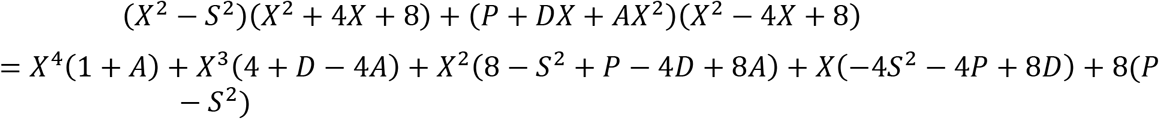

With critical feedback, this should have the same roots as:

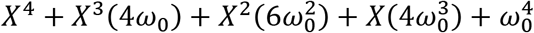

We numerically solve the system of equations:

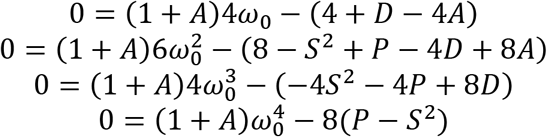

The resulting time to recover balance and critical gains as a function of S are shown in Fig 10.D-C in green.

#### g) Integral feedback

We consider that the system has integral feedback *i* such that:

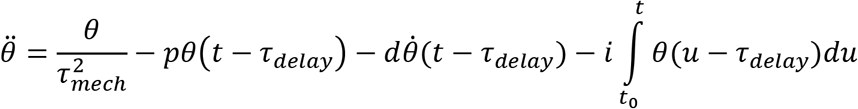

Introducing the dimensionless variable 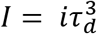, the characteristic equation is given by:

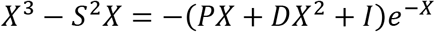

Using the first order Pade approximation, the characteristic equation becomes:

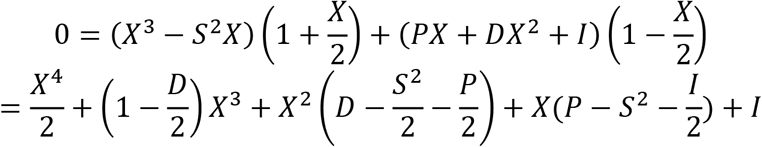

With critical feedback, this should have the same roots as:

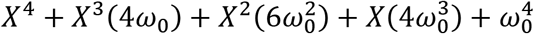

We numerically solve the system of equations:

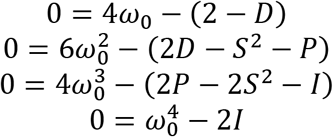

#### h) Multi-dimensional systems

We consider an N-dimensional dynamical system with state **θ** and delayed feedback control **C**, whose dynamics are governed by:

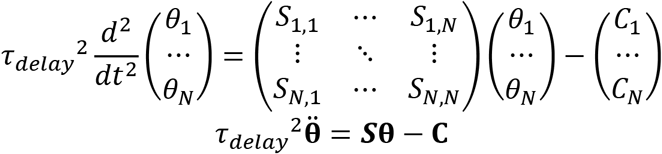

We consider the generic case, in which the transpose ***S***^*T*^ of the dynamics matrix ***S*** is diagonalizable, and introduce the basis set (***e***_1_,…,***e_N_***) of eigenvectors of ***S***^*T*^ and their corresponding eigenvalues (*s*_1_,…,*s_N_*), such that for every *i*:

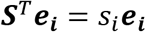

We use this basis set to perform a transformation of coordinates of the state **θ** into ***α***, such that, for every *i, α_i_* is the dot product of the vectors ***e_i_*** and **θ**:

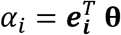

Each component *α_i_* follows the dynamical equation:

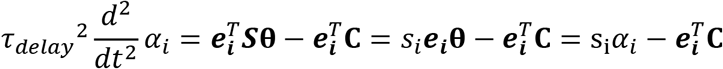

The dynamics is thus decomposed into a set of N components, each of which follows a single-dimensional dynamics for which the analysis presented in the previous sections holds. When considering a model of human stance with several joints (such as the ankle, knee and hip), the dynamics can be decomposed into such a set of components. Each of these components is a linear combination of the various angles, and its dynamics is governed by a mechanical time constant *τ_i_* such that 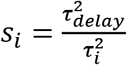.

