## Supplementary Text for "Anticipatory coadaptation of ankle stiffness and sensorimotor gain for standing balance"

Charlotte Le Mouel, Romain Brette

#### I. Stability analysis

The system is a linearized inverted pendulum, with an external forcing $F$:

$$\tau_{delay}{}^2\ddot{\theta} - S\,\theta = F \tag{1}$$

The system is controlled with delayed proportional-derivative feedback $u$, based on the observed value of $\theta$, written $\theta_{obs}$:

$$u = (G\theta_{obs} + D\tau_{delay}\dot{\theta}_{obs})(t - \tau_{delay}) \tag{2}$$

To determine the stability of the system, we analyse how perturbation signals propagate through the system. we consider 2 types of perturbation: a perturbation δ in the external force, and noise η in the observation process:

$$F = -u + \delta$$
$$\theta_{obs} = \theta + \eta$$

The block diagram of the controlled system with noise is shown in Figure S1.A.

Replacing in (1) and (2):

$$\tau_{delay}{}^2\ddot{\theta} - S\,\theta = \delta - G\theta(t-\tau) - D\tau_{delay}\,\dot{\theta}(t-\tau) - G\eta(t-\tau) - D\tau_{delay}\,\dot{\eta}(t-\tau)$$
$$\tau_{delay}{}^2\ddot{\theta} - S\,\theta + G\theta(t-\tau) + D\tau_{delay}\,\dot{\theta}(t-\tau) = \delta - G\eta(t-\tau) - \tau_{delay}D\,\dot{\eta}(t-\tau) \tag{3}$$

The unforced motion of the system (for $\delta = 0$ and $\eta = 0$) is given by the solutions to the homogeneous equation:

$$\tau_{delay}{}^2\ddot{\theta} - S\,\theta + G\theta(t-\tau) + D\tau_{delay}\,\dot{\theta}(t-\tau) = 0$$

The solutions to this equation are called the modes of the system. After an arbitrary perturbation, the unforced motion of the system is a weighted sum of such modes.

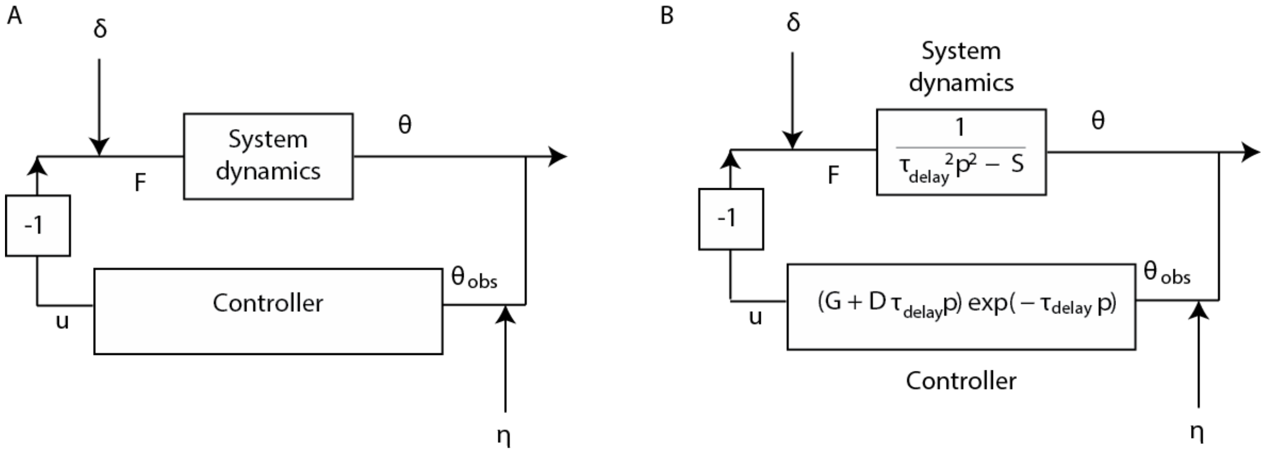


*Figure S1 Block diagram of the controlled system*

*A. Noise is injected into the system both at the level of the motor command (δ) and at the level of the sensory feedback (η). B. Transfer function of the system dynamics and controller.*

### 1) Propagation of exponential signals and derivation of the characteristic equation

Since the system is linear, we only need to consider the response to exponential signals $e^{pt}$ where $p$ is a complex number. The response to a sum of exponential signals is then the sum of the responses to each exponential signal. We therefore consider perturbations of the form:

$$\delta(\mathrm{t}) = \delta_0 e^{pt}$$
$$\eta(\mathrm{t}) = \eta_0 e^{pt}$$

Then the response of the system is also an exponential signal such that:

$$\theta(\mathrm{t}) = \theta_0 e^{pt}$$

The transfer function of the system dynamics and controller are shown in Figure S1.B.

Replacing in (4):

$$\left(\tau_{delay}{}^2 p^2 - S + Ge^{-p\tau_{delay}} + D\tau_{delay} p e^{-p\tau_{delay}}\right)\theta_0 e^{pt} = \delta_0 e^{pt} - \left(G + D\tau_{delay} p\right) e^{-p\tau_{delay}} \eta_0 e^{pt}$$

The solutions to the characteristic equation $D(p) = 0$ correspond to the modes of the homogeneous equation, where:

$$D(p) = \tau_{delay}{}^2 p^2 - S + Ge^{-p\tau_{delay}} + D\tau_{delay} p e^{-p\tau_{delay}}$$

If there exists $p$ with positive real part such that $D(p) = 0$, then the system is unstable. Indeed, the amplitude of an exponential signal is $e^{Re(p)t}$, thus if $Re(p) > 0$ the amplitude of the mode grows exponentially with time. If this mode is excited by a perturbation at one point, then even after the end of the perturbation, the system will diverge.

This can also be seen by looking at the transfer function of the system:

$$\theta_0 = \frac{\delta_0 - (G + D\tau_{delay} p) e^{-p\tau_{delay}} \eta_0}{\tau_{delay}{}^2 p^2 - S + (G + D\tau_{delay} p) e^{-p\tau_{delay}}}$$

If $D(p) = 0$, then the response of the system to a perturbation $e^{pt}$ diverges, since the denominator of the transfer function becomes zero. The roots of the denominator are called the poles. The system is therefore stable if and only if its transfer function has no poles with positive real part (Aström & Murray, 2010).

Note that the denominator is the same whether noise in injected into the motor or the sensory process. The system is therefore either robust to both sensory and motor noise or robust to neither. Indeed, stability only depends on the behaviour of the homogeneous equation.

The difficulty in assessing stability comes from the feedback delay, which introduces the $e^{-p\tau_{delay}}$ term: because of this term, the characteristic equation has an infinite number of roots, and there is no straightforward criterion to determine stability (Michiels & Niculescu, 2007). Consider for example, the characteristic equation $1 + e^{-p\tau_{delay}} = 0$. For all integers $n$, $p = \frac{i(1+2n)\pi}{\tau_{delay}}$ is a root of the equation: the equation therefore has an infinite number of roots.

To assess stability, we will therefore use the Nyquist criterion, introduced by Nyquist (Nyquist, 1932) and described in the following section. For convenience, we will apply the Nyquist to the open-loop transfer function:

$$OL(p) = \frac{\left(G + D\tau_{delay} p\right) e^{-\tau_{delay} p}}{\tau_{delay}{}^2 p^2 - S}$$

The transfer function of the system is related to the open loop transfer function according to:

$$\theta_0 = \frac{\frac{\delta_0}{\tau_{delay}{}^2 p^2 - S} - OL(p)\eta_0}{1 + OL(p)}$$

The poles of the transfer function are therefore the zeros of $1 + OL(p)$.

### 2) Nyquist criterion

We therefore seek to determine whether $f(p) = 1 + OL(p)$ has zeros with positive real part.

For this, we will use Cauchy's residue theorem, which states that the integral of a function $g(p)$ (which must be analytical except at a number of poles and zeros) over a contour in the complex plane is equal to the sum of the residues of $g(p)$ at each of its poles and zeros within the region encompassed by that contour.

We first introduce the function $g(p) = \frac{f\prime(p)}{f(p)}$ whose sum of residues within a region is equal to the difference between the number of zeros and poles of $f(p)$ within that region. we then integrate this function over the Nyquist contour which encompasses the right half-plane. we thus determine the number of zeros of $f(p)$ with positive real part.

#### a. Residues of g(p)

The function $g(p) = \frac{f\prime(p)}{f(p)}$ is analytical except at the poles and zeros of $f(p)$.

We therefore use Cauchy's residue theorem, which states that the integral of $g(p)$ over a contour $\Gamma$ is equal to the sum of the residues of $g(p)$ at each of its poles and zeros within this contour (which correspond to the poles and zeros of $f(p)$).

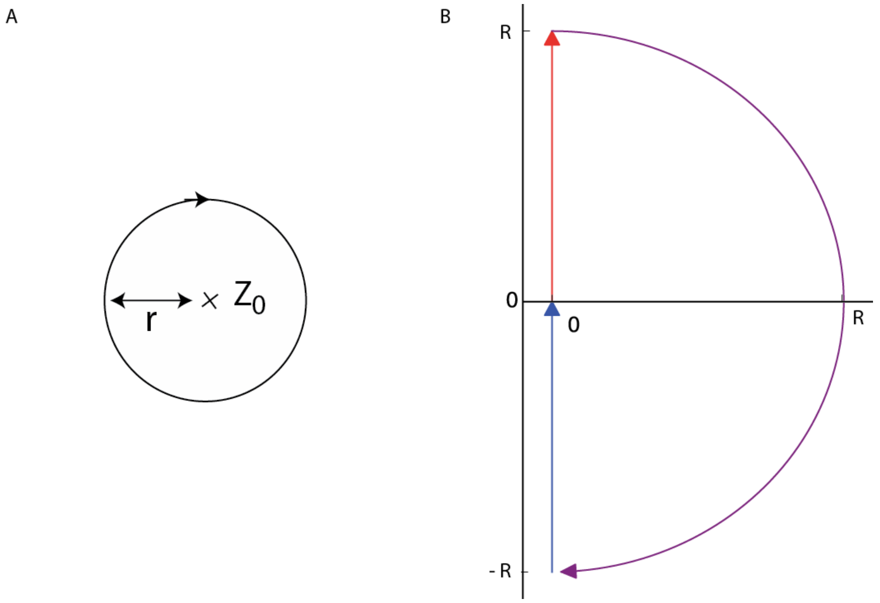


*Figure S2 Calculation of residuals within the Nyquist contour*

*A. Neighbourhood of a zero or pole B. D-shaped contour of a given radius R*

*Zeros*

We consider $z_0$ a zero of $f(p)$ of multiplicity m, and write:

$$f(p) = (p - z_0)^m h(p)$$

$$g(p) = \frac{f'(p)}{f(p)} = \frac{m(p - z_0)^{m-1} h(p)}{(p - z_0)^m h(p)} + \frac{(p - z_0)^m h'(p)}{(p - z_0)^m h(p)}$$

$$= \frac{m}{p - z_0} + \frac{h'(p)}{h(p)}$$

There exists a neighbourhood of $z_0$ for which $h(p)$ and therefore $\frac{h\prime(p)}{h(p)}$ is analytic. We choose $r$ such that the circle $\Gamma_{r,z_0}$ centered at $z_0$ and of radius $r$ is contained within this neighbourhood (Figure S2.A). The residue of $g(p)$ at $z_0$ is equal to the integral of $g(p)$ around the circle:

$$Res(z_0) = \oint_{\Gamma_{r,z_0}} g(p)dp = \oint_{\Gamma_{r,z_0}} \frac{m}{p - z_0} dp + \oint_{\Gamma_{r,z_0}} \frac{h'(p)}{h(p)} dp$$

Since $\frac{h\prime(p)}{h(p)}$ is analytic within this circle:

$$\oint_{\Gamma_{r,z_0}} \frac{h'(p)}{h(p)} dp = 0$$

We introduce the change of variables: $p = z_0 + re^{i\phi}, dp = ire^{i\phi}d\phi$

$$\oint_{\Gamma_{r,z_0}} \frac{m}{p - z_0} dp = m \int_{\phi=0}^{2\,\pi} \frac{ire^{i\phi}d\phi}{re^{i\phi}} = 2i\,\pi\,m$$

*Poles*

We consider $p_0$ a pole of $f(p)$ of multiplicity n, and write:

$$f(p) = \frac{k(p)}{(p - p_0)^n}$$

$$f'^{(p)} = \frac{k'(p)}{(p - p_0)^n} + k(p)(-n)\frac{1}{(p - p_0)^{n+1}}$$

$$g(p) = \frac{f'(p)}{f(p)} = \frac{k'(p)}{k(p)} - \frac{n}{p - p_0}$$

There exists a neighbourhood of $p_0$ for which $k(p)$ is analytic. The residue of $g(p)$ at $p_0$ is equal to the integral of $g(p)$ around the circle $\Gamma_{r,p_0}$ centred on $p_0$ and included within this neighbourhood:

$$\oint_{\Gamma_{r,p_0}} \frac{-n}{p - p_0} dp = -2\,i\,\pi\,n$$

Therefore, the integral of $g(p)$ over the a contour is equal to $2i\,\pi\,(m-n)$, where $m$ is the number of zeros of $f(p)$ and $n$ is the number of poles of $f(p)$ within that contour.

##### b. Nyquist contour

Since the region we am interested in is the entire right half-plane (the region of the complex plane with positive real part), we will use the Nyquist contour $\Gamma_N$, which is the limit, for $R \to +\infty$, of the D-shaped contour (traversed counterclockwise) defined by:

- $i\omega$ for $\omega$ ranging from $-R$ to $+R$
- $R\,e^{i\,\phi}$ for $\phi$ ranging from π/2 to - π/2

This contour is plotted in Figure S2.B for a given radius $R$, and for $R \to +\infty$ this contour encompasses all of the complex plane with positive real part. Note: we use the convention that clockwise curves are oriented positively.

##### c. Geometrical interpretation

$$\oint_{\Gamma_N} g(p)dp = \oint_{\Gamma_N} \frac{f'(p)}{f(p)}dp = \oint_{\Gamma_N} \frac{d}{dp}\log\big(f(p)\big)\,dp = \oint_{\Gamma_N} \frac{d}{dp}\log(|f(p)|)\,dp + \oint_{\Gamma_N} \frac{d}{dp} i\arg\big(f(p)\big)\,dp$$

The variation of $|f(p)|$ over the closed contour $\Gamma_N$ is zero.
The number of zeros minus poles is therefore equal to the winding number $w$:

$$w = \frac{1}{2\,\pi} \oint_{\Gamma_N} \frac{d}{dp}\arg\big(f(p)\big)\,dp$$

Geometrically, $w$ corresponds to the number of clockwise loops effected by $f(p)$ around 0 when $p$ ranges over the Nyquist curve.

The Nyquist criterion thus states that a system described by the open-loop transfer function $OL(p)$ with $n$ poles with positive real part is stable if and only if the curve described by $OL(p)$, as $p$ ranges over the Nyquist contour, loops $n$ times counter-clockwise around the point $-1$.

#### 3) Application to our system

The open-loop transfer function is given by:

$$OL(p) = \frac{\big(G + D\tau_{delay}p\big)e^{-\tau_{delay}p}}{\tau_{delay}{}^2p^2 - S}$$

$f(p) = 1 + OL(p)$ has a unique pole with positive real part $p = \sqrt{S}/\tau_{delay}$. Therefore, the system is stable if and only if the integral of $g(p)$ over the Nyquist contour is equal to $-2i\,\pi$. Thus, for the system to be stable, the open-loop transfer function $OL(p)$ must loop once counterclockwise around the point $-1$ when $p$ ranges over the Nyquist contour.

The second part of the Nyquist contour, defined by $R\,e^{i\,\phi}$ for $\phi$ ranging from π/2 to - π/2, with $R \to +\infty$, maps onto the point 0. Indeed:

$$\lim_{R\to+\infty} OL\big(R\,e^{i\,\phi}\big) = 0$$

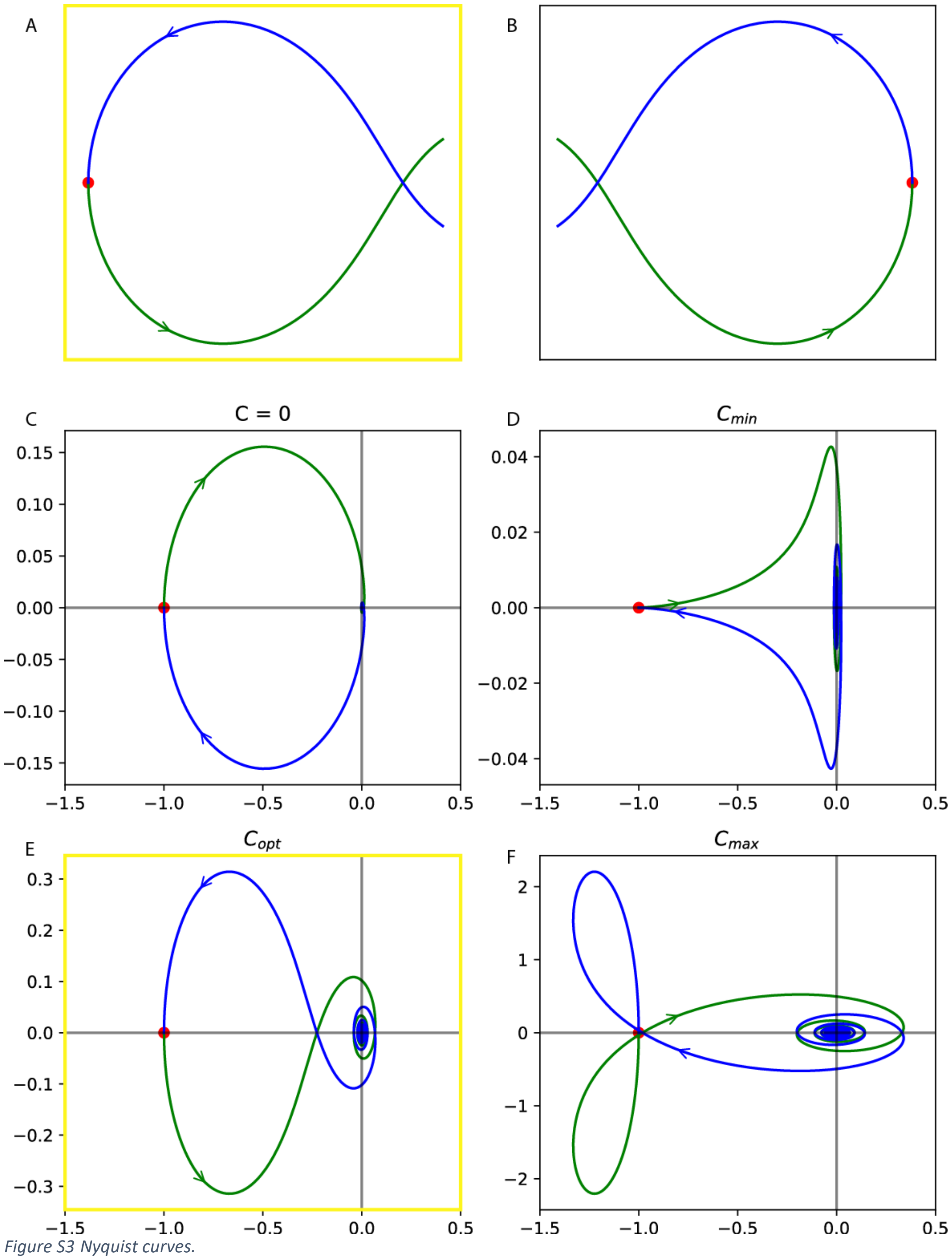


*Figure S3 Nyquist curves.*

*The Nyquist curve is plotted in blue for X < 0, in green for X > 0, and in red for X = 0. Two possibilities for a counterclockwise loop are schematically illustrated in A. and B. The Nyquist curve is plotted for C = 0 in panel C., minimal damping C = 1 in panel D., critical damping $C_{opt}$ in panel E., and maximal damping $C_{max}$ in panel F. The only curve in panels C-F which contains a counterclockwise loop is the one in panel E, which corresponds to the situation in panel A.*

The first part of the Nyquist contour, defined by $i\omega$ for $\omega$ ranging from $-\infty$ to $+\infty$, maps onto the curve described by:

$$OL(i\omega) = \frac{\left(G + D\tau_{delay} i\omega\right)e^{-\tau_{delay} i\omega}}{-\tau_{delay}{}^2\omega^2 - S}$$

We introduce the dimensionless parameter:

$$X = \tau_{delay}\omega$$

Then the Nyquist curve can equivalently be described by (with $X$ ranging from $-\infty$ to $+\infty$):

$$OL(X) = -\frac{(G + iDX)e^{-iX}}{X^2 + S}$$

To disentangle the effects of $G$, $D$ and $S$ on stability, we introduce $C = \frac{D}{G}$ such that the gain $G$ simply scales the curve defined by:

$$OL(X) = -G\frac{(1 + iCX)e^{-iX}}{X^2 + S}$$

We first determine the parameters $(S, C)$ for which there exists a counterclockwise loop in the curve. For clarity, in figures we use the gain $G = S$. Then, for a given set of admissible $(S, C)$, we determine the minimal and maximal gains for which the curve loops around $-1$.

#### a. $(S, C)$ parameters with a counter-clockwise loop

Description of the curve:

- The curve for $X < 0$ is the symmetric with respect to the real axis of the curve for $X > 0$
- For $X = \pm\infty$, $OL = 0$ because of the $X^2$ in the denominator
- For $X = 0$, $OL = -G/S$

We consider the first intersection point of the Nyquist curve with the real axis for $X > 0$, and denote it $X_{int}(C, S)$. There are 2 options for a counter-clockwise loop, schematically illustrated in Figure S3:

A. $-G/S < OL(X_{int}(C, S))$ and the imaginary part is negative for $X \in [0, X_{int}(C, S)]$ (Figure S3.A)
B. $-G/S > OL(X_{int}(C, S))$ and the imaginary part is positive for $X \in [0, X_{int}(C, S)]$ (Figure S3.B)

*Determination of the intersection point*

We expand the open-loop into real and imaginary parts:

$$OL(X) = -G\frac{(1 + iCX)(\cos(X) - i\sin(X))}{X^2 + S} = \frac{G}{X^2 + S}(1 + iCX)(-\cos(X) + i\sin(X))$$

$$= \frac{G}{X^2 + S}\left(-\left(\cos(X) + CX\sin(X)\right) + i\left(-CX\cos(X) + \sin(X)\right)\right)$$

The sign of the imaginary part is thus the same as the sign of:

$$I(C, X) = -CX\cos(X) + \sin(X)$$

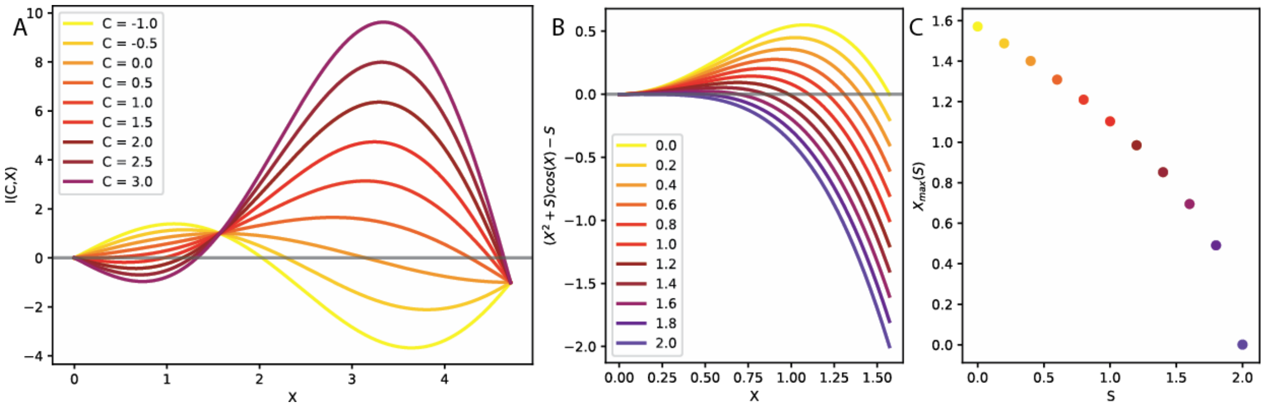


*Figure S4 Determination of the intersection point.*

*A. The curve I(C,X) is illustrated as a function of X for different values of C ranging from -1 to 3. B. The curve $f_S(X)$ is illustrated as a function of X for different values of S ranging from 0 to 2. C. The maximal value of the intersection point X for which there exists a counter-clockwise loop in the Nyquist curve is illustrated as a function of S.*

This function is illustrated in Figure S4.A for different values of $C$. The intersection point therefore depends only on $C$ and not on S, and we write it $X_{int}(C)$. It satisfies :

$$\frac{\tan(X_{int}(C))}{X_{int}(C)} = C$$

As can be seen in Figure S4.A:

- For $C < 1$, $X_{int}(C) \in ]\pi/2, 3\pi/2[$ and $I(C,X) > 0$ for $X \in [0, X_{int}(C)]$
- For $C > 1$, $X_{int}(C) \in ]0, \pi/2[$ and $I(C,X) < 0$ for $X \in [0, X_{int}(C)]$

Indeed, a first order expansion around $X = 0$ gives:

$$I(C,X) \propto -CX + X = X(1-C)$$

We now consider the value of the open-loop function at the intersection point:

$$OL(X_{int}) = \frac{-G}{{X_{int}}^2 + S}(\cos(X_{int}) + CX_{int}\sin(X_{int})) = \frac{-G}{{X_{int}}^2 + S}\frac{1}{\cos(X_{int})}(\cos(X_{int})^2 + \sin(X_{int})^2)$$
$$= \frac{-G}{({X_{int}}^2 + S)\cos(X_{int})}$$

*Stability requires C > 1*

As we have shown, for $C < 1$, $X_{int}(D) \in ]\pi/2, 3\pi/2[$ therefore $\cos(X_{int}) < 0$ and $OL(X_{int}) > 0 > -G/S$. Moreover, for $C < 1$, the imaginary part is positive for $X \in [0, X_{int}(D)]$. Thus, the Nyquist curve satisfies neither conditions A. nor conditions B., and the system is not stable: there is no counter-clockwise loop in the curve. The Nyquist curves for $C = 0$ and $C = 1$ are illustrated in Figure S3 (respectively panel C. and D.).

*Stability requires $S < 2$*

As we have shown, for $C > 1$, the imaginary part is negative for $X \in [0, X_{int}(C)]$. Stability therefore requires satisfying condition A., ie:

$$-\frac{G}{S} < OL\big(X_{int}(C)\big) = \frac{-G}{(X_{int}(C)^2 + S)\cos\big(X_{int}(C)\big)}$$
$$0 < (X_{int}(C)^2 + S)\cos\big(X_{int}(C)\big) - S$$

$X_{int}(C)$ is an increasing function of $C$, ranging from 0 for $C \to 1$ to $\frac{\pi}{2}$ for $C \to +\infty$.

We therefore define $f_S: X \in \left[0, \frac{\pi}{2}\right] \to (X^2 + S)\cos(X) - S$

$$f_S(0) = 0$$

$$f_S\left(\frac{\pi}{2}\right) = -S$$

This function is illustrated in Figure S4.B for different values of S.

The derivative is given by:

$$f_S'(X) = 2X\cos(X) - S\sin(X)$$

For $S \geq 2$, it is negative throughout the range $\left[0, \frac{\pi}{2}\right]$, therefore $f_S(X)$ is also negative throughout this range (Figure S4.B), and there exists no value of $C$ for which the system is stable.

*Stability requires* $C < C_{max}(S)$

For $S < 2$, $f_S'(0) > 0$, therefore there exists a range of values $X \in [0, X_{max}(S)]$ for which $f_S(X) > 0$. The value of $X_{max}(S)$ for different values of S is shown in Figure S4.C. Thus there exists a range of $C \in [1, C_{max}(S)]$ for which the Nyquist curve has a counterclockwise loop. $C_{max}(S)$ is given by:

$$C_{max}(S) = \frac{\tan(X_{max}(S))}{X_{max}(S)}$$

The Nyquist curve for the critical damping $C_{opt} > 1$ (as derived in the following section V.2) and the maximal value of damping $C_{max}(S)$ are illustrated in Figure S3 (respectively panel E. and F.)

##### b. Feedback gain $G$ for which the loop encompasses $-1$

Finally, for a given value of speed and damping, the range of gains which can stabilize the system is given by:

$$-\frac{G}{S} < -1 < \frac{-G}{(X_{int}(C)^2 + S)\cos(X_{int}(C))}$$

$$S < G < (X_{int}(C)^2 + S)\cos(X_{int}(C))$$

The minimal gain is thus $G = S$ for all values of $C$.

The maximal gain depends on $C$, and follows a curve parametrized by $X \in [0, X_{max}(S)]$ :

$$G = (X^2 + S)\cos(X)$$

$$C = \frac{\tan(X)}{X}$$

Note that $G(0) = S$ and $G(X_{max}(S)) = S$.

#### 4) Simulations

##### a. Feedback gain $P$

The response of systems with various feedback gains is shown in Figure S5.A for the relative speed $S = 0.1$ and the critical damping $D_{crit}$ for that relative speed:

- For $P < S$ (dashed red line), the feedback is not strong enough to prevent falling, and the system is unstable.
- For $P = S$ (dashed black line), the feedback is just strong enough to prevent falling, but not strong enough to bring the system back to its initial position: this is the lower limit of stability.
- For $P = G_{max}$ (full black line), the feedback elicits oscillations whose amplitude neither increases nor decreases with time: this is the upper limit of stability.

For gains between $S$ and $G_{max}$ (blue and green dashed and full lines), the system is stable:

- For $P = P_{crit}$ the critical gain (full green line) the perturbation is cancelled the fastest without oscillations.
- For $P < P_{crit}$ (dashed blue line), the perturbation is cancelled more slowly.

– For $P > P_{crit}$ (full blue line), there are oscillations.

#### b. Feedback damping $D$

The response of systems with various feedback dampings is shown in Figure S5.B, for the relative speed $S = 0.1$ and the critical gain $P_{crit}$. If the damping is too low (dashed red and black lines), slow oscillations appear, whereas if it is too large (full black line), fast oscillations appear. For intermediate values of damping (full and dashed blue and green lines), the perturbation is cancelled the fastest without oscillations for $D = D_{crit}$ (full green line).

#### c. Relative speed $S$

The response of systems with various relative speeds and critical feedback parameters is shown in Figure S5.C:

– As $S$ approaches 2 (blue line), an initial perturbation of amplitude 1 is amplified 400 times before it is cancelled by the feedback (note the difference in the scale of the y axis between panel C and panels A and B).

– For $S = 2$ (black line), the amplitude of the oscillations neither increases nor decreases with time: this is the upper limit of stability.

– For $S > 2$ (red line), no feedback gains are able to stabilize the system, and the amplitude of the oscillations grows with time.

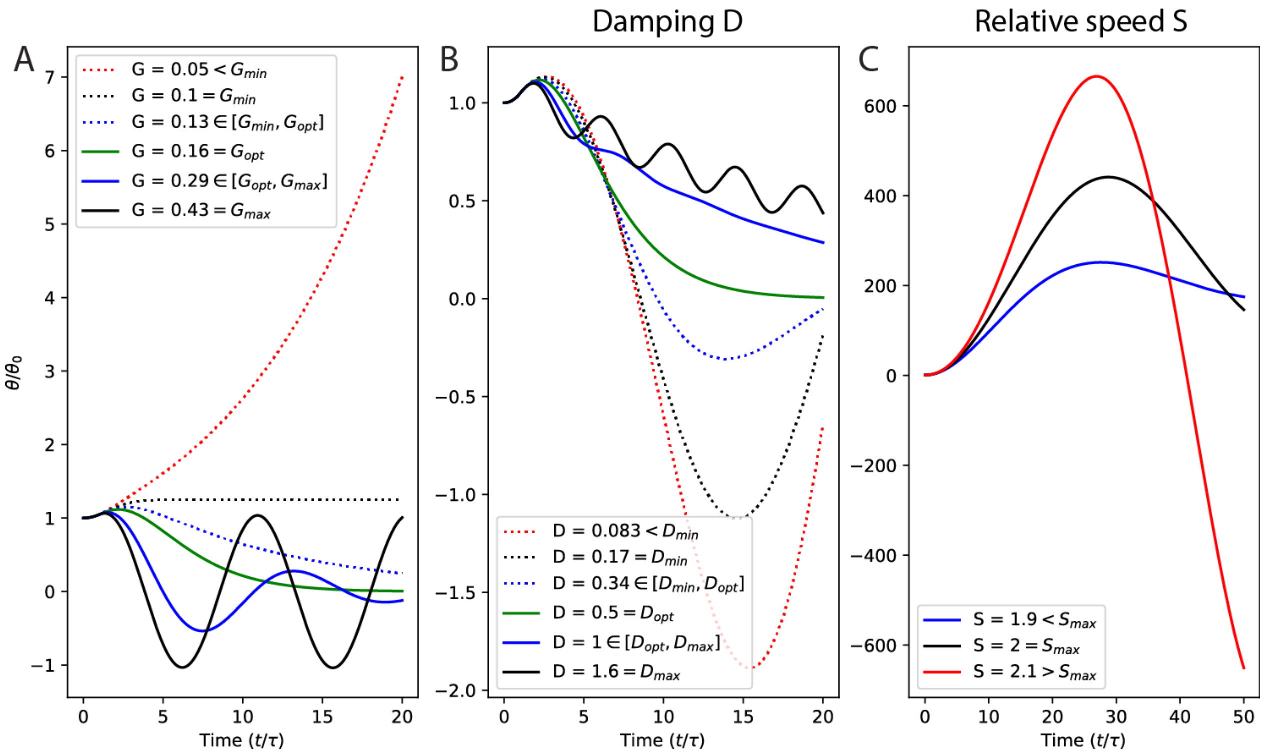


*Figure S5 Response to a perturbation.*

*The position ϑ of the system to a perturbation of arbitrary magnitude ϑ_0 is displayed as a function of time (normalized to the response delay τ_delay). A. Response of a system with relative speed S = 0.1, damping Dopt (S), and various gains. B. Response of a system with relative speed S = 0.1, gain Gopt (S), and various dampings. C. Response of the system with various relative speeds, and critical gain and damping Gopt (S),Dopt (S). Stable systems are in blue and green, unstable systems in red, and systems at the border of stability in black. Systems with feedback parameters lower than the critical values are in dashed lines.*

### II. Pade approximation

The first order Pade approximation of the delay is given by:

$$e^{-X} = \frac{e^{-X/2}}{e^{X/2}} \approx \frac{1 - X/2}{1 + X/2}$$

The Pade approximation consists in approximating the function $\theta(t - \tau_{delay})$ by a function $\theta_{approx}(t)$ which follows:

$$\frac{\tau_{delay}}{2}\dot{\theta}_{approx}(t) + \theta_{approx}(t) = \theta(t) - \frac{\tau_{delay}}{2}\dot{\theta}(t)$$

Examples:

1. Suppose $\theta(t)$ is a step function defined by (Figure S6.A in black):

- $\theta(t) = 0 \; for \; t < 0$
- $\theta(t) = 1 \; for \; t > 0$

Then $\theta_{approx}(t) = 1 - \exp(-\frac{2t}{\tau_{delay}})$ (Figure S6.A in red).

2. Suppose $\theta(t)$ is a sinusoidal function defined by $\theta(t) = \exp(i\omega t)$ (Figure S6.B, C, in black), then:

$$\theta_{approx}(t) = \frac{1 - \frac{i\omega\tau_{delay}}{2}}{1 + \frac{i\omega\tau_{delay}}{2}} \exp(i\omega t)$$

The amplitude is:

$$\sqrt{\frac{(1 + (\frac{\omega\tau_{delay}}{2})^2)^2}{(1 + (\frac{\omega\tau_{delay}}{2})^2)^2}} = 1$$

The phase lag is:

$$\phi\left(\frac{1 - \frac{i\omega\tau_{delay}}{2}}{1 + \frac{i\omega\tau_{delay}}{2}}\right) = -2\arctan\left(\frac{\omega\tau_{delay}}{2}\right)$$

This corresponds to a time lag of $\tau_{delay}$ for small $\omega\tau_{delay}$ (Figure S6.C in red) and π for large $\omega\tau_{delay}$ (Figure S6.B in red).

Thus, the Pade approximation $\theta_{approx}(t)$ corresponds to $\theta(t - \tau_{delay})$ if $\theta$ changes slowly compared to $\tau_{delay}$ (Figure S6.C), whereas fast variations in $\theta$ are distorted (Figure S6.A, B).

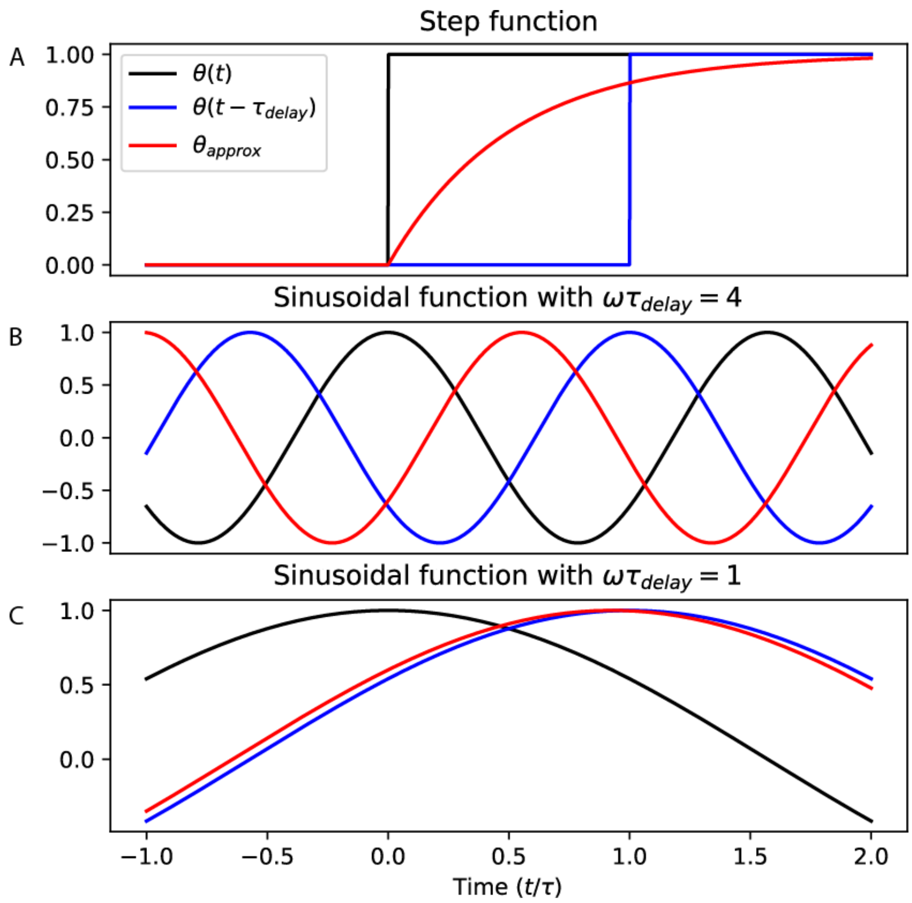


*Figure S6 Pade approximation of the time delay.*

*A. Step function in black, delayed step function in blue and Pade approximation of the delayed step function in red. B-C Sinusoidal function with $\omega\tau_{delay} = 4$ and 1 (respectively for panels B and C) in black, delayed sinusoidal function in blue and Pade approximation of the delayed sinusoidal function in red.*
